# Plasticity in astrocyte subpopulations regulates heroin relapse

**DOI:** 10.1101/2020.07.22.216036

**Authors:** Anna Kruyer, Ariana Angelis, Constanza Garcia-Keller, Hong Li, Peter W. Kalivas

**Author notes:** Correspondence should be addressed to A.K. (173 Ashley Ave, MSC510, Charleston, SC, 29425).

## Abstract

Opioid use disorder (OUD) produces detrimental personal and societal consequences. Astrocytes are a major cell group in the brain that receives little attention in mediating OUD. We determined how astrocytes and the astroglial glutamate transporter, GLT-1, in the nucleus accumbens core adapt and contribute to heroin seeking in rats. Seeking heroin, but not sucrose, produced two transient forms of plasticity in different astroglial subpopulations. Increased morphological proximity to synapses occurred in one subpopulation and increased extrasynaptic GLT-1 expression in another. Augmented synapse proximity by astroglia occurred selectively at D2-dopamine receptor expressing dendrites, while changes in GLT-1 were not neuron-subtype specific. Importantly, mRNA-antisense inhibition of either morphological or GLT-1 plasticity promoted cue-induced heroin seeking. We show that heroin cues induce two distinct forms of transient plasticity in separate astroglial subpopulations that dampen heroin relapse.

**TEASER:** Different subpopulations of astrocytes engage with accumbens synapses to dampen heroin relapse.

## INTRODUCTION

Relapse to opioid use is a leading cause of death in the United States. Decades of research demonstrate the importance of glutamate dysregulation at nucleus accumbens core (NAcore) synapses as a causative factor in relapse-like behavior in animal models (*1*). Glutamate dysregulation in animal models of addiction is due in large part to changes in NAcore astroglia that express the glutamate transporter GLT-1, which terminates the actions of the bulk of synaptically-released glutamate (*2, 3*). Addictive substances, including alcohol, nicotine, psychostimulants, and opioids induce an enduring downregulation of GLT-1 on astrocytes in the NAcore (*4*). Furthermore, astroglial processes that insulate synapses and take up glutamate during synaptic transmission retract from NAcore synapses after extinction from cocaine, heroin, and methamphetamine use, but not following sucrose self-administration and extinction (*5–7*). Synaptic retraction of astrocyte processes and downregulation of GLT-1 disrupt glutamate homeostasis at NAcore synapses, permitting spillover of synaptic glutamate and postsynaptic potentiation required for drug-associated cues to initiate drug seeking (*8*).

We previously found that synaptic retraction of NAcore astroglia after heroin withdrawal is partially and transiently reversed during cue-induced heroin seeking (*6*). We hypothesize that heroin cue-induced increased proximity of astroglia to synapses increases the synaptic adjacency of GLT-1 on perisynaptic astroglial processes, thereby serving to minimize spillover of synaptically-released glutamate that mediates cue-induced drug seeking. We also sought to determine whether synaptic reassociation of astroglial processes stimulated by heroin-associated cues was selective for either of the two main neural subtypes in the NAcore, D1 or D2 receptor-expressing medium spiny neurons (D1- or D2-MSNs) (*9*), which exert opposing control over drug-seeking behaviors (*10–13*).

We used confocal microscopy to measure the synaptic co-registration of labeled astrocytes and astroglial GLT-1 in the NAcore after extinction or reinstatement of heroin or sucrose seeking. We found that cued reinstatement of heroin, but not sucrose, seeking produced two transient adaptations that parsed into distinct astroglial subpopulations. In one subpopulation, heroin cues increased synaptic adjacency of astroglial processes without increasing surface-proximal GLT-1, and in the second subpopulation, cued reinstatement elevated surface GLT-1 on astroglia situated more than 250 nm from the synaptic cleft. We independently disrupted the two forms cue-induced plasticity in each subpopulation of NAcore astroglia by knocking down GLT-1 or by reducing synthesis of ezrin, an astroglial selective actin binding protein that mediates morphological plasticity in distal astroglial processes (*14–18*). Either knockdown augmented the capacity of cues to induce heroin seeking. When we assessed whether astrocytes exhibited a bias in their synaptic adjacency with D1- or D2-MSN synapses, we found that astrocytes selectively associated with D1-MSN synapses after extinction of heroin seeking and retracted from D2-MSNs, and that this pattern was reversed during cue-reinstated heroin seeking. Moreover, the cue-induced increase in surface GLT-1 was not detected adjacent to either D1- or D2-MSN dendrites during reinstated heroin seeking, supporting a functional role for GLT-1-deficient astroglial processes in shaping synaptic activity. In keeping with abundant literature demonstrating suppression of synaptic activity by perisynaptic astroglia (*19, 20*), we propose that astrocyte insulation of D1-MSN synapses after extinction training suppresses D1-MSN activity to reduce seeking, while retraction from D2-MSNs permits their potentiation. The reversal of this pattern during cue presentation would instead permit D1-MSN potentiation, which is known to drive cued drug seeking (*10*). In all, our data show that astrocyte morphological plasticity is neuron subtype-selective and that two distinct forms of astroglial plasticity are transiently induced by heroin cues in separate subpopulations of astrocytes to dampen cue-induced heroin seeking.

## RESULTS

### GLT-1 surface expression was transiently elevated during cued heroin seeking

To examine whether enhanced synaptic proximity of NAcore astroglia suppressed cue-induced heroin seeking through changes in synaptic adjacency of GLT-1, NAcore astroglia were selectively labeled with a membrane-bound fluorescent reporter prior to operant training. Rats were trained to self-administer heroin or sucrose and reward delivery was paired with light and tone cues (Fig. 1A). Importantly, heroin intake was equivalent in rats placed in the extinction, 15-min or 120-min reinstated groups (Fig. 1A inset). Animals that received yoked saline delivery and cues served as controls for heroin-trained rats and animals that received yoked cues were controls for sucrose-trained rats. Operant responding was extinguished by removing both the cues and reinforcers, and a portion of rats were reinstated by restoring conditioned cues without reinforcers to active lever pressing for 15 or 120 min (Fig. 1B). Rats used to generate the data in Figs. 1A-B were included in a previous study (*6*) and in the present report mCherry transduced astrocytes in tissue slices from these rats were immuno-labeled for both Synapsin I and GLT-1. NAcore slices from each treatment group (yoked, extinguished, 15-min reinstated, 120-min reinstated) were double-labeled for GLT-1 and the presynaptic marker Synapsin I and imaged using confocal microscopy (Fig. 1C). Total GLT-1, co-registered Synapsin I, and co-registered GLT-1 and Synapsin I were quantified and normalized to the volume of each mCherry-labeled astrocyte (Fig. 1D-E). To estimate the proportion of surface-proximal GLT-1, we digitally isolated GLT-1 within 250 nm of the astroglial membrane as a proportion of total GLT-1 in each astrocyte (Fig. 1F-H).

**Fig. 1.**
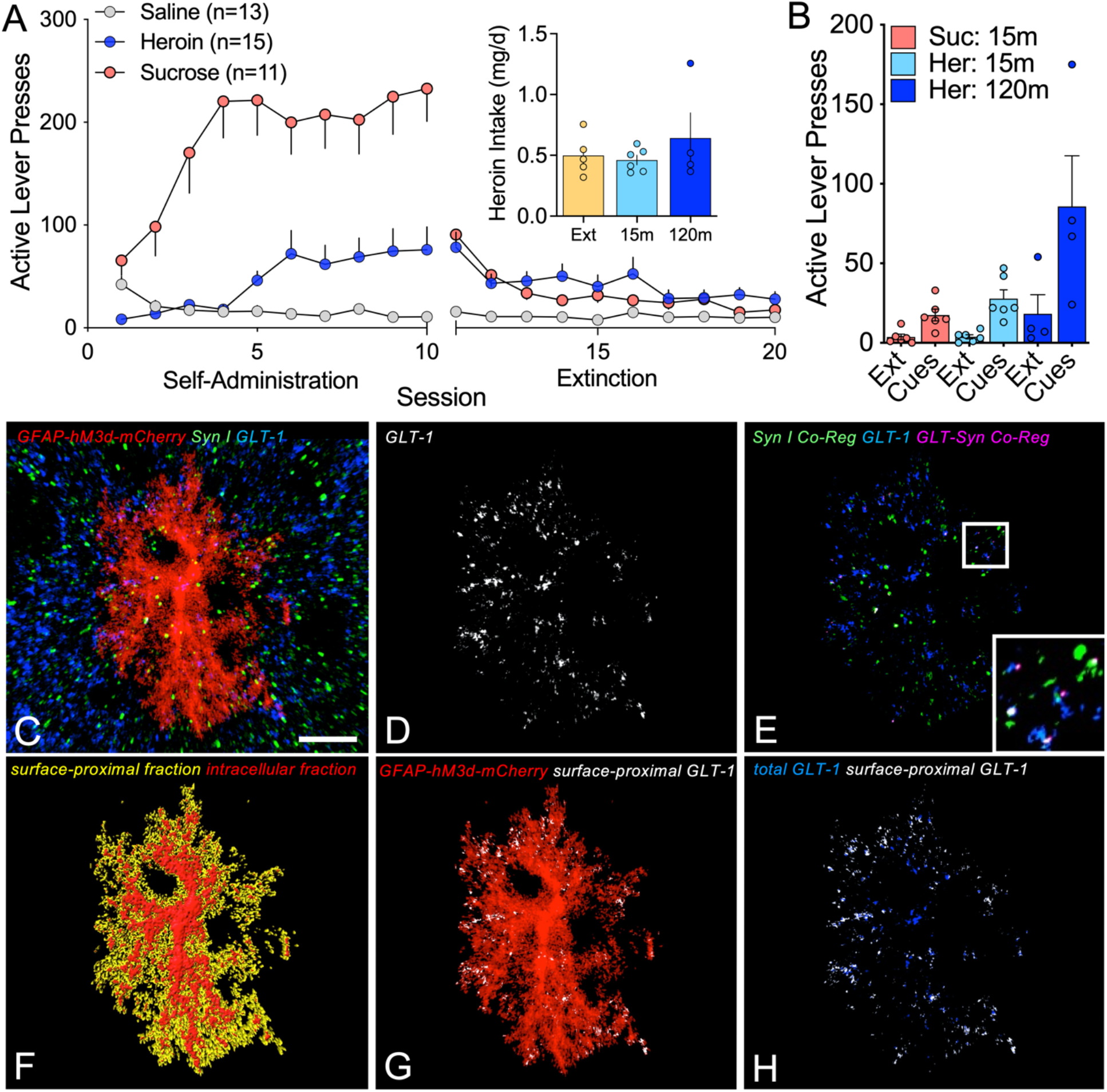
Workflow used for confocal analysis of astroglial morphology and surface-proximal GLT-1. (**A**) Rats were trained to self-administer heroin or sucrose over 10 consecutive days of operant training and reward delivery was paired with light and tone cues. During extinction training, heroin or sucrose and cues were not delivered in response to active lever pressing and operant responding gradually decreased. Heroin- and sucrose-trained rats differed in active lever pressing during self-administration (2-way ANOVA, F_1,124_=33.44, p<0.0001). Inset shows that groups of heroin-trained animals took similar amounts of heroin (1-way ANOVA F_2,12_=0.9187, p=0.4984). (**B**) 24h after the last extinction session, reinstated animals were placed in the operant chamber and cues were restored to the active lever for 15- or 120-min to reinstate seeking (2-way ANOVA Session F_1,26_=16.07, p=0.0005). (**C**) Z-series depicting an NAcore astrocyte transfected with AAV5/GFAP-hM3d-mCherry (red) and immuno-labeled for Synapsin I (green) and GLT-1 (blue). (**D**) GLT-1 immunoreactivity co-registered with mCherry in (**C**) is shown in white. (**E**) Co-registration of GLT-1 (blue) with Synapsin I (green) from the region occupied by the astrocyte in (**C**) is shown in pink. (**F**) Digital rendering of the astroglial surface (yellow) was used to identify GLT-1 signal within 250 nm of the cell membrane (**G**-**H**, white) relative to total GLT-1 from the same astrocyte (**H**, blue). Bar in (**C**)= 10 μm. In (**A**, inset), Ext, extinguished; 15m, 15-min cued reinstatement; 120m, 120-min cued reinstatement. In (**B**), Suc, sucrose; Her, heroin; Ext, extinction.

Co-registration of the astroglial membrane with Synapsin I was reduced after withdrawal from heroin (Fig. 2A), but not sucrose (Fig. S1A). We also found reductions in total astrocyte GLT-1 expression after extinction from heroin (Fig. 2B), but not sucrose self-administration (Fig. S1B). Although total GLT-1 was reduced in heroin extinguished rats, the proportion of surface-proximal GLT-1 was unaltered in extinguished rats (Fig. 2C). Moreover, 15 min of cue exposure transiently increased the proportion of astroglia expressing the highest levels of surface-proximal GLT-1 (Fig. 2D). Both the increase in synapse-proximal astroglia and GLT-1 after 15-min of heroin cue were transient and returned to extinction levels by 120-min after initiating cued reinstatement (Fig. 2A,C-D). Interestingly, the ratio of surface:total GLT-1 was reduced in the NAcore of rats extinguished from sucrose (Fig. S1C), but not altered by cued reinstatement (Fig. S1C-D). The co-registration of GLT-1 with Synapsin I was reduced after extinction from heroin self-administration (Fig. 2E), but not sucrose self-administration (Fig. S1E). Despite increases in surface GLT-1 and synaptic adjacency by NAcore astroglia, the proximity of GLT-1 to the presynaptic marker Synapsin I was not restored by cued heroin seeking (Fig. 2E-F). Instead, the increase in surface-proximal GLT-1 in reinstated animals was targeted extrasynaptically (>250 nm from the synapse) (Fig. 2G-H). In sucrose-trained rats, levels of synaptic and extrasynaptic GLT-1 were not changed by extinction training or cued reinstatement of sucrose seeking (Fig. S1F-H). Importantly, extinction from heroin self-administration did not change total Synapsin I expression in the NAcore (Fig. S2).

**Fig. 2.**
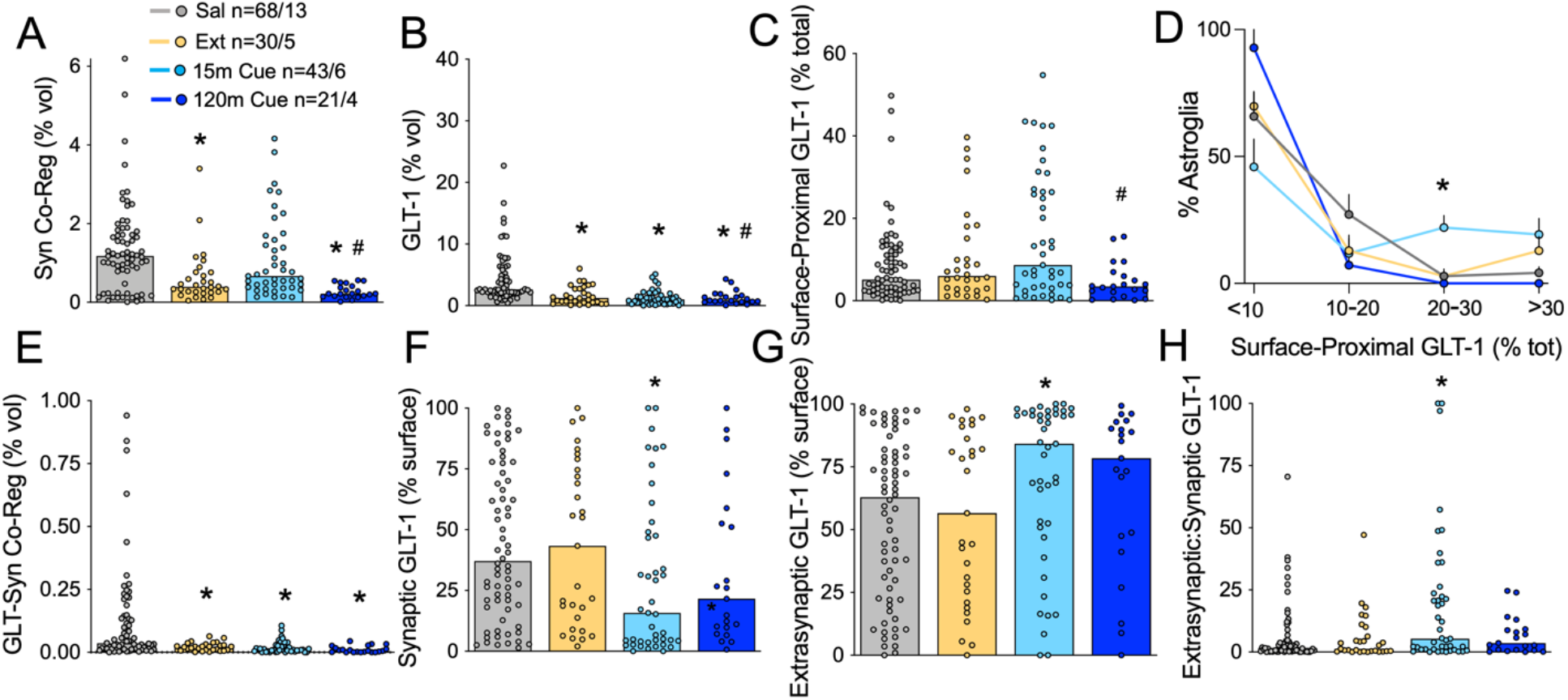
Extrasynaptic GLT-1 was transiently elevated during heroin seeking. We previously found that co-registration of labeled NAcore astroglia with Synapsin I is reduced during extinction from heroin (**A**, Kruskal-Wallis=30.17, p<0.0001) and that 15-min of cued heroin seeking restores synaptic insulation by astrocytes. (**B**) Withdrawal from heroin self-administration produced a downregulation of GLT-1 on NAcore astroglia, whether or not rats were reinstated (Kruskal-Wallis=51.77, p<0.0001). (**C**) Surface-proximal GLT-1, shown here as percent of total GLT-1 from each astrocyte, was increased during active seeking (15m Cue) when compared to a timepoint at which cues no longer evoke seeking (i.e. cue extinction; 120m Cue, Kruskal-Wallis=9.848, p<0.05). A greater proportion of astrocytes exhibited high levels of surface-proximal GLT-1 after 15-min of heroin cues compared to yoked saline controls (**D**, 2-way ANOVA, F_6,48_=2.904, p<0.05). Co-registration of GLT-1 with the presynaptic marker Synapsin I was found to be reduced in heroin-trained rats, and was not restored during cued reinstatement (**E**, Kruskal-Wallis=35.48, p<0.0001). When synaptic and extrasynaptic fractions of surface GLT-1 were analyzed separately, we found that 15-min of cued heroin seeking decreased synaptic GLT-1 (**F**, Kruskal-Wallis=9.493, p<0.05) and increased extrasynaptic GLT-1 (**G**, Kruskal-Wallis=9.493, p<0.05). The ratio of extrasynaptic and synaptic GLT-1 illustrates the robust increase in extrasynaptic GLT-1 during cued heroin seeking (**H**, Kruskal-Wallis=9.476). N shown in (**A**) as cells/animals. *p<0.05 compared to yoked control, #p<0.05 compared to 15-min reinstated using Dunn’s test. Sal, yoked saline; Ext, extinguished; 15m Cue, 15-min cued reinstatement; 120m Cue, 120-min cued reinstatement.

### Astrocyte heterogeneity in heroin cue-induced plasticity

The fact that astrocyte synaptic proximity was transiently increased by 15 min of cued heroin reinstatement, but the increase in surface-proximal GLT-1 did not co-register with Synapsin I indicates that these cue-induced astroglial adaptations may occur in distinct subpopulations of astrocytes. Indeed, principal component analysis (PCA, Fig. S3) (*21*) of astroglia from saline-, heroin- and sucrose-trained rats identified three distinct clusters of astroglia we arbitrarily refer to as types 1-3: type 1, astrocytes with high synaptic co-registration; type 2, astrocytes with high levels of extrasynaptic GLT-1, and type 3, astrocytes with low-to-moderate synaptic adjacency and surface GLT-1 expression (Fig. 3A). A majority of NAcore astrocytes from yoked saline rats were in the type 1 and 3 subpopulations (Fig. 3B, E). Following extinction from heroin self-administration there was a loss of type 1 and an increase in type 2 astrocytes (Fig. 3C, E). Fifteen min of cued heroin reinstatement resulted in a transient restoration of type 1 astrocytes, and an increase in the proportion of type 2 astroglia (Fig. 3D, E). In contrast with heroin self-administration and akin to saline rats, all three treatment groups of sucrose-trained rats contained predominately type 1 and 3 astrocytes (Fig. S4A). Along with the lack of GLT-1 co-registration with Synapsin I during cued reinstatement (Fig. 2E), these data are consistent with the presence of three distinct astroglial populations in heroin-seeking animals, which is also indicated by the lack of GLT-1 immuno-staining in some NAcore astroglia (Fig. S5). Fig. 3F illustrates the distinct morphology and GLT-1 localization characteristic of each astroglial type.

**Fig. 3.**
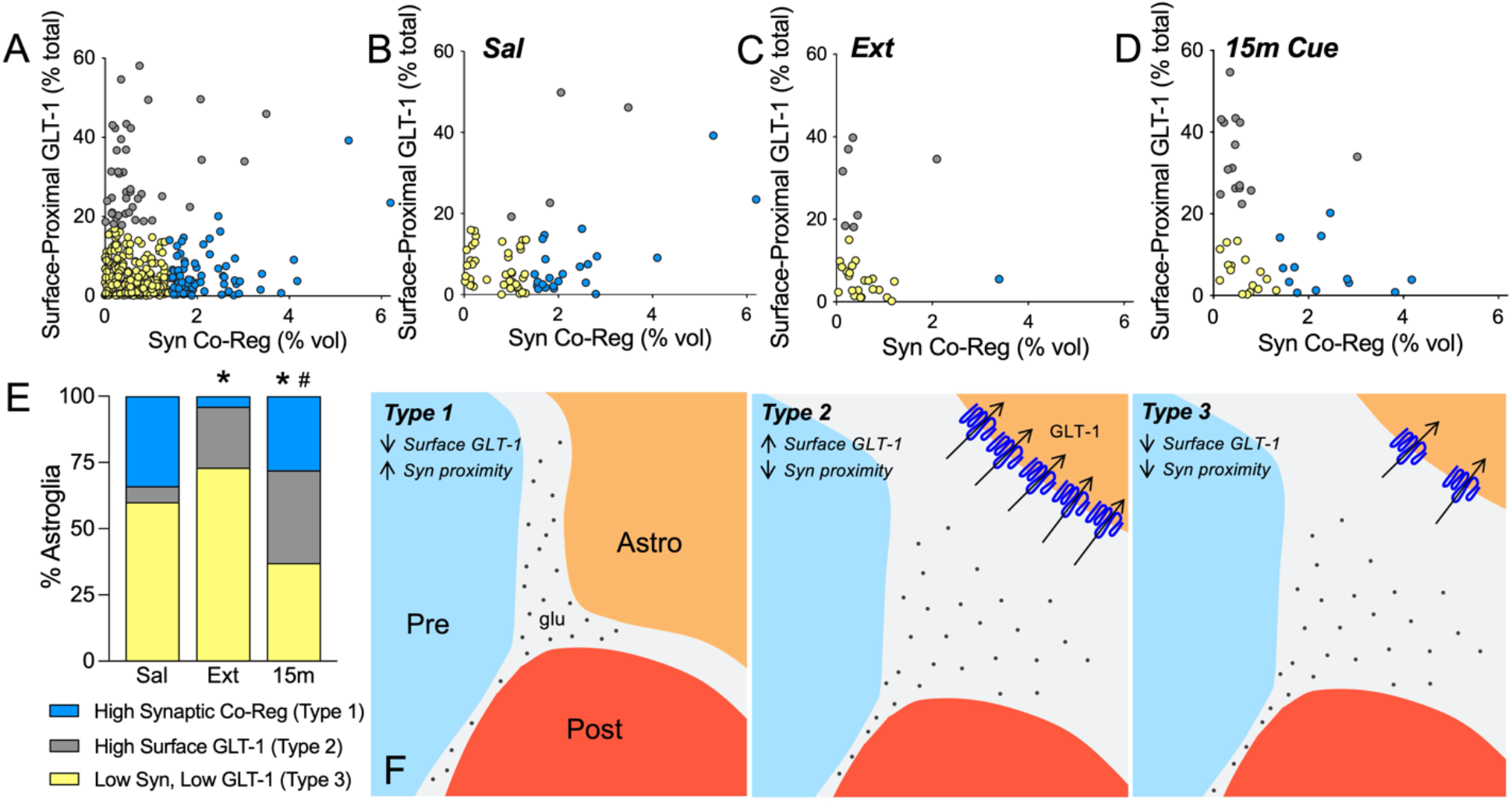
Cue-induced increases in astrocyte motility and GLT-1 surface expression occurred in different astroglial subpopulations. (**A**) Principal component analysis was used to identify subpopulations of NAcore astroglia according to their synaptic adjacency and levels of surface-proximal GLT-1. Astrocytes with high synaptic adjacency (blue) or high levels of surface-proximal GLT-1 (gray) were identified as separate cell clusters. Distribution of clusters in tissue from yoked saline, extinguished or cue-reinstated rats is shown in (**B**-**D**). Astrocytes with high surface-proximal GLT-1, but low synaptic co-registration were largely absent from yoked control rats, but emerged after operant heroin training (**E**, Chi^2^=14.10 *p=0.0018 Ext. vs. Sal). Half of astroglia in cue-reinstated rats exhibited one or the other type of transient plasticity (**E**, Chi^2^=15.97, *p=0.0006 vs. Sal; Chi^2^=11.20, #p=0.0074 vs. Ext). (**F**) Schematic illustrating 3 astroglial subtypes identified by PCA. Type 1 astroglia expressed low levels of GLT-1, but exhibited high measures of synaptic-adjacency (**F**, left panel). Type 2 astroglia expressed high levels of largely extrasynaptic GLT-1 (blue) and were observed heroin training, but were not abundant in control animals (**F**, middle panel). Type 3 astroglia were predominant in control animals and had low-to-moderate GLT-1 expression and synaptic adjacency (**F**, right panel).

### G_q_ signaling in NAcore astroglia increased synaptic adjacency, but not surface GLT-1

G_q_-coupled signaling through metabotropic glutamate receptor mGluR5 on cultured astroglia promotes ezrin-dependent astrocyte fine process motility (*15*). Moreover, cue-induced heroin seeking is associated with increased ezrin phosphorylation (*6*) and astrocyte proximity to synapses (Fig. 2A), and increases in synaptic glutamate spillover in NAcore that occur during cued reinstatement of drug seeking (*22*) could stimulate mGluR5 on astroglial processes (*23*). To determine if G_q_-type mGluR signaling triggers cue-induced astrocyte process motility, a G_q_-coupled designer receptor activated by designer drug (DREADD) was delivered selectively to NAcore astroglia (Fig. 4A). Rats were trained to self-administer heroin before undergoing 10 days of extinction training (Fig. 4B). In lieu of reinstatement, animals were given clozapine N-oxide (CNO) or vehicle and their brains were removed after 30-min for analysis of astrocyte synaptic adjacency and surface-proximal GLT-1 expression. CNO delivery increased the co-registration of the astroglial membrane with Synapsin I-positive puncta (Fig. 4C), but did not impact surface-proximal GLT-1 levels (Fig. 4D). These data indicate that G_q_ signaling in NAcore astroglia triggers morphological plasticity during reinstatement, while the increase in extrasynaptic GLT-1 expression (Fig. 2G-H) likely involves a different signaling cascade.

**Fig. 4.**
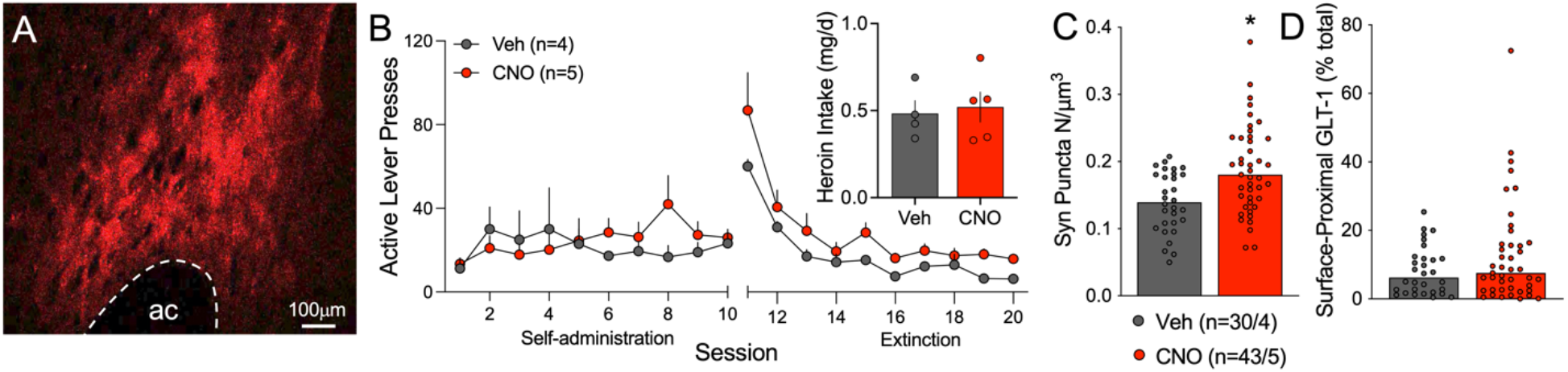
G_q_-signaling increased synaptic adjacency in NAcore astroglia from heroin-trained rats. After viral delivery of G_q_-DREADD in NAcore astroglia (**A**, red), rats were trained to self-administer heroin before undergoing extinction training (**B**). The following day, rats received i.p. injections of vehicle or CNO, but did not undergo cued reinstatement. Heroin intake did not differ between vehicle- or CNO-treated rats (**B**, inset, t_7_=0.173, p=0.7602). (**C**) G_q_ signaling increased co-registration of NAcore astroglia with near-adjacent immunolabeled Synapsin I puncta (Mann-Whitney U=363, p=0.0013), but did not impact levels of surface-proximal GLT-1 (**D**, Mann-Whitney U=568, p=0.3934). In (**A**), ac, anterior commissure. In (**B**-**C**), Veh, Vehicle. In (**C**), N shown as cells/animals.

### Reducing cue-induced plasticity in astroglial subpopulations increased heroin seeking

To determine whether cue-induced type 1 and type 2 astroglial plasticity (as shown in Fig. 3F) impacted heroin seeking, we implanted cannulae above the NAcore and trained rats to self-administer heroin (Fig. 5A). Rats were divided to maintain equal heroin intake across the reinstated groups (Fig. 5B) and received bilateral infusions of an inert control vivo-morpholino antisense oligomer, or an oligo targeted to GLT-1 (*24*) or ezrin (*6*) on days 6-8 of extinction training. Rats were reinstated by exposure to heroin cues and either ezrin or GLT-1 knockdown potentiated cued seeking (Fig. 5C). Thus, the presence of either type 1 or type 2 astroglial plasticity during cued heroin seeking serves a compensatory function to suppress heroin seeking.

**Fig. 5.**
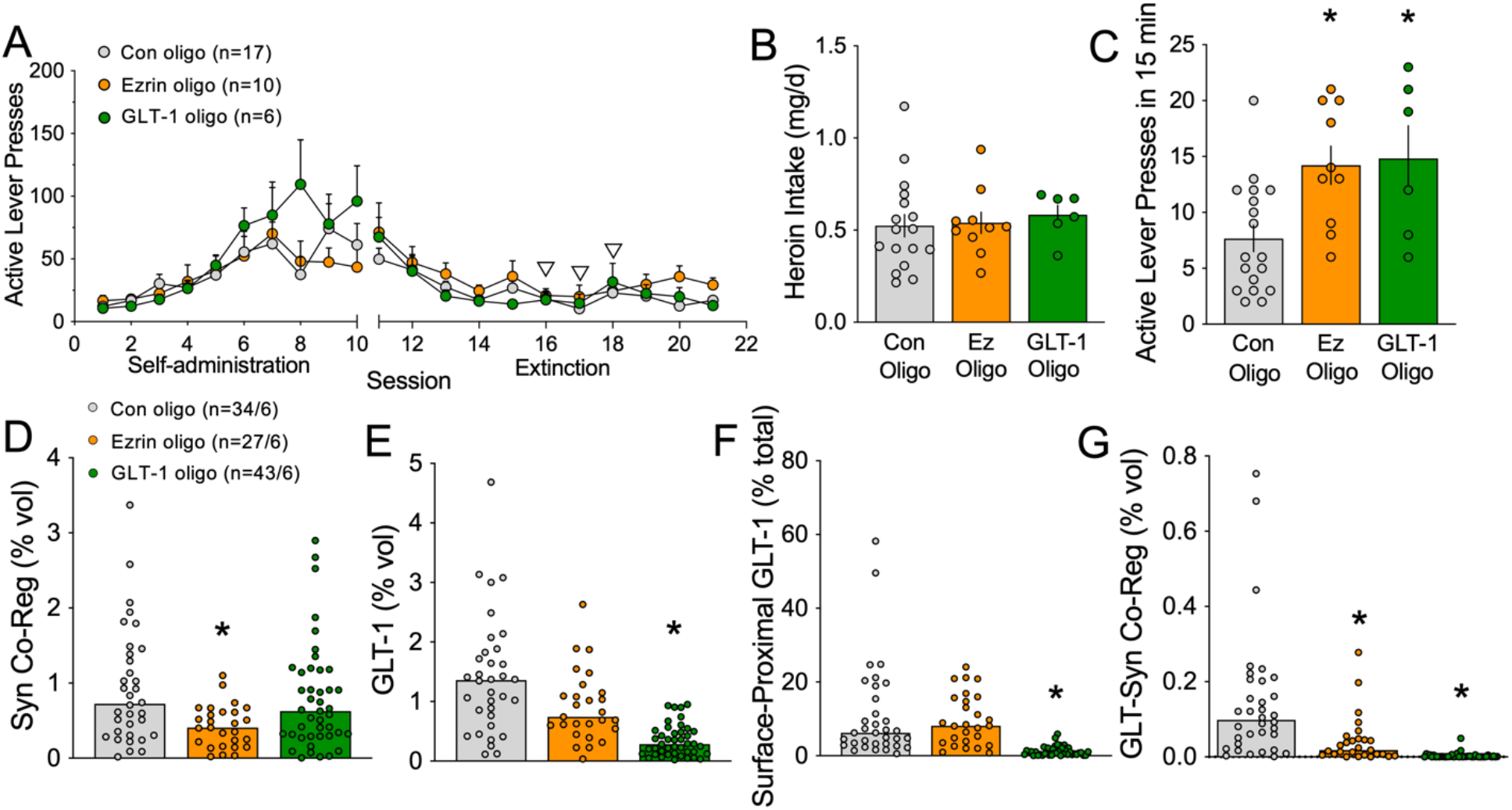
Impairing cue-induced astrocyte motility or surface GLT-1 elevated heroin seeking. (**A**) Rats were trained to self-administer heroin over 10 consecutive days. Starting on day 6 of extinction training for 3 consecutive days (white arrowheads), animals received NAcore infusions of an ezrin antisense oligomer, a GLT-1 antisense oligomer or a control oligomer. Animals in each treatment group did not differ in total heroin intake during self-administration. (**B**) Rats receiving different oligo treatments did not differ in heroin intake (1-way ANOVA F_2,30_=0.1739, p=0.8412). 24h after the final extinction session, animals were reinstated for 15-min by exposure to light and tone cues previously paired with heroin delivery. (**C**) Rats that underwent ezrin or GLT-1 knockdown pressed higher on the active lever during the 15-min reinstatement session (1-way ANOVA F_2,30_=6.230, p=0.0055, *p=0.0058 Con vs. Ezrin Oligo, *p=0.0104 Con vs. GLT-1 Oligo using Fisher’s test). (**D**) Astrocytes from rats treated with the ezrin oligo exhibited a significant reduction in synaptic co-registration, consistent with knockdown of astrocyte peripheral process motility (Kruskal-Wallis=15.85, p=0.0004, *p=0.0002 vs. Con). The GLT-1 oligo did not impact synaptic co-registration by NAcore astroglia (p=0.4676 vs. Con). (**E**) GLT-1 levels were unchanged by ezrin antisense oligo delivery (p=0.3888), but were reduced by treatment with the GLT-1 oligo (Kruskal-Wallis=54.85, p<0.0001, *p<0.0001). (**F**) Likewise, surface-proximal GLT-1 was reduced after application of the GLT-1 oligo (Kruskal-Wallis=80.22, p<0.0001, *p<0.0001 vs. Con), but not the ezrin oligo (p>0.9999 vs. Con). (**G**) The co-registration between GLT-1 and Synapsin I on astroglia was significantly reduced after ezrin (Kruskal-Wallis=87.13, p<0.0001, *p=0.0147) or GLT-1 knockdown (*p<0.0001). In (**A**-**D**), Con, control oligo; Ez, ezrin oligo; GLT-1, GLT-1 oligo. In (**D**), N shown as cells/animals.

Confirming that cannulae placement and morpholino infusions did not impact cue-induced astrocyte plasticity, both type 1 and type 2 adaptations were observed during 15 min of cue-induced heroin seeking in rats treated with control oligo (compare Fig. S4B with Fig. 3D). Ezrin knockdown reduced the synaptic proximity of astroglia in the NAcore (Fig. 5D) and GLT-1 knockdown reduced total and surface proximal GLT-1 (Fig. 5E-F). Validating the specificity of the oligos and supporting the presence of distinct subpopulations of astrocytes, the ezrin-targeted oligo selectively eliminated type 1 astrocytes while the GLT-1 targeted oligo selectively eliminated type 2 astrocytes in rats reinstated by heroin cues for 15 min (Fig. S4B). Moreover, GLT-1 knockdown did not alter synaptic proximity of NAcore astroglia (Fig. 5D), and ezrin knockdown did not impact levels of total GLT-1 or surface-proximal GLT-1 (Fig. 5E-F). As expected, independent knockdown of either GLT-1 or ezrin reduced GLT-1 co-registration with Synapsin I (Fig. 5G).

Taken together these data show that all three types of astroglia are present during cue-induced heroin seeking (Fig. 3F). Akin to extinguished rats, heroin cues are associated with type 2 astroglia having increased surface-proximal expression of GLT-1, and akin to their presence in control saline or sucrose rats, cues induce the presence of type 1 astroglia that have high synaptic proximity. The subpopulations are distinct cells and are regulated by distinct signaling pathways. Importantly, the presence of either type 1 or type 2 subpopulations serves to inhibit cue-induced heroin seeking.

### NAcore astrocytes differentially associated with D1- and D2-MSNs at baseline and after heroin

Activity in the two main neuronal subtypes in the NAcore, D1- and D2-MSNs promotes drug seeking and extinction of seeking, respectively (*25, 26*) and individual astrocytes in the dorsal striatum signal uniquely with these two neuronal subclasses (*19, 27*). We hypothesized that the different functional types of astroglia depicted in Fig. 3F would associate uniquely with D1- and D2-MSNs, thereby contributing to their differential functional roles in heroin extinction and cue-induced seeking. To test this hypothesis, we labeled astrocytes and neurons in male and female D1- and D2-Cre transgenic rats before training animals to self-administer heroin for 10 days (Fig. 6A-B). Some rats were then reinstated for 15-min by exposure to heroin-associated cues (Fig. 6C). Isolated astroglia were imaged and their co-registration with virally-labeled D1- or D2-MSN dendrites and the synaptic marker Synapsin I or GLT-1 was quantified (Fig. 6D-G). Pooling data from D1- and D2-Cre rats revealed the same synaptic retraction by astroglia from NAcore dendrites after extinction and re-insertion of astroglial processes toward synapses during cued reinstatement (Fig. 6H) as was observed in Sprague-Dawley rats (Fig. 2A). Interestingly, when astrocyte-synapse co-registration data were analyzed separately in D1- and D2-Cre rats, the astrocyte association with D1-MSN synapses was lower in yoked saline animals compared with D2-MSN synapses (compare saline-treated groups in Fig. 6I-J). After extinction training, astrocyte co-registration with Synapsin I was increased at D1-MSN synapses (Fig. 6I) and decreased at D2-MSN synapses (Fig. 6J). Fifteen minutes of exposure to heroin cues restored astrocyte association with both synaptic types to control levels (Fig. 6I-J).

**Fig. 6.**
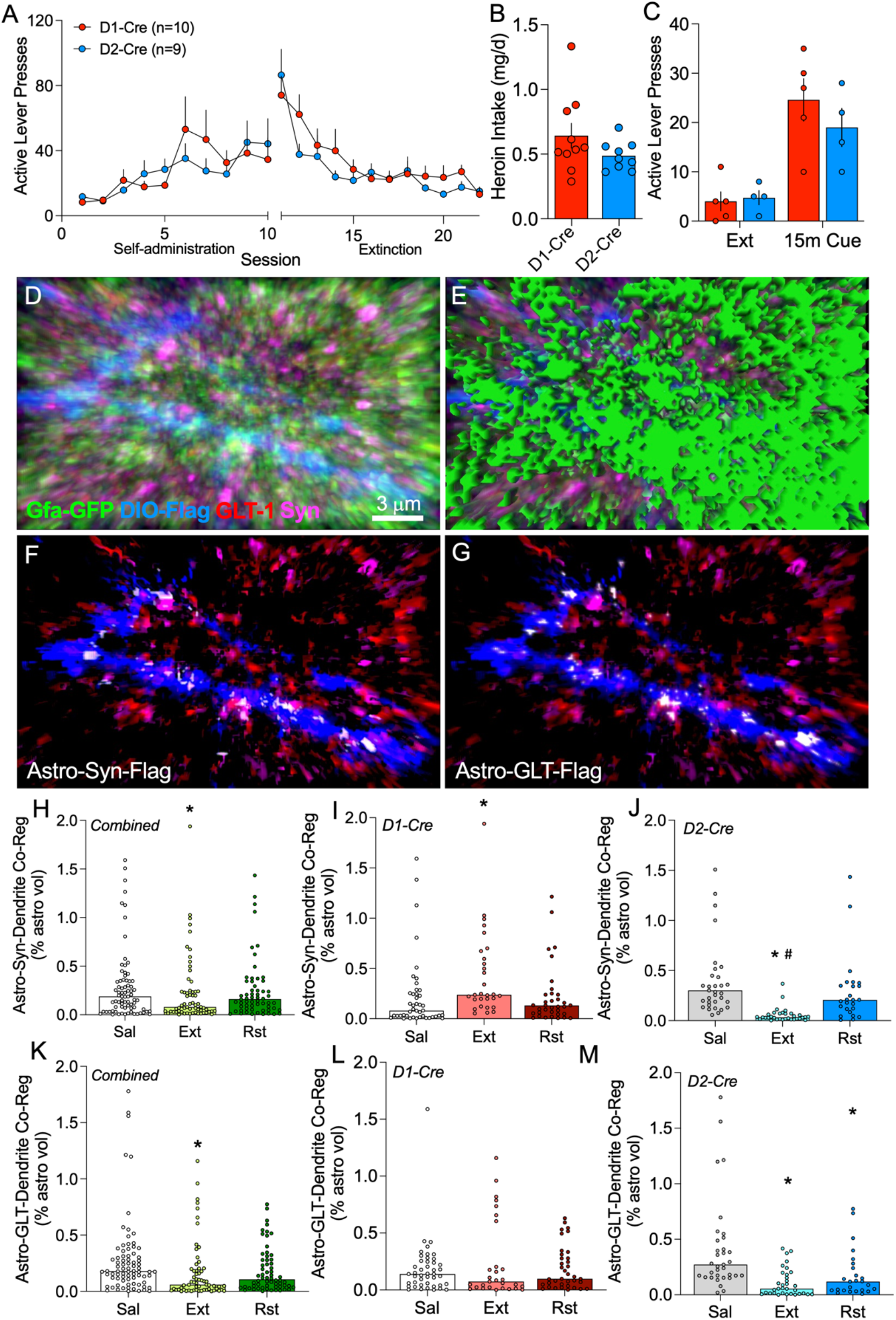
Astrocytes increased their adjacency to D1-MSN synapses but retracted from D2-MSN synapses after extinction training. (**A**) Male and female D1- and D2-Cre rats were trained to self-administer heroin over 10 days before undergoing extinction training. (**B**) Heroin intake did not differ between the strains (t_17_=1.449, p=0.1656). (**C**) 24h after the last extinction session, lever pressing was reinstated by exposure to heroin-conditioned cues for 15-min (2-way ANOVA Time F_1,7_=20.36 p=0.0028, Genotype F_1,7_=0.9944 p=0.3519). (**D**) NAcore tissue from these animals was immunolabeled to identify D1- or D2-MSN dendrites (blue), GLT-1 (red), and Synapsin I (pink). Signal associated with NAcore astroglia (green, **D**-**E**) was isolated to quantify the degree of triple co-registration between astroglia and D1- or D2-MSN synapses (**F**, white), or astroglial GLT-1 and D1- or D2-MSN dendrites (**G**, white). Synaptic association by astroglia was reduced by extinction training after heroin self-administration when quantified in a cell-nonspecific manner (**H**, Kruskal-Wallis=4.617 p=0.0994, *p=0.0380 vs. Sal). When analyzed separately, astrocyte association with D1-MSN synapses was low at baseline and was increased by extinction training (**I**, Kruskal-Wallis=12.35 p=0.0021, *p=0.0014 vs. Sal). Instead, D2-MSN synapses had high astrocyte insulation at baseline (p<0.0001 vs. D1-Cre, Sal using Dunn’s test) that was reduced by extinction training and recovered during cued reinstatement (**J**, Kruskal-Wallis=45.84 p<0.0001, *p<0.0001 vs. Sal, #p<0.0001 vs. Rst). Surface-proximal, dendrite-associated GLT-1 was decreased overall after extinction training in NAcore astrocytes, and was restored during 15-min cued reinstatement (**K**, Kruskal-Wallis=15.23 p=0.0005, *p<0.05 vs. Sal). However, the increase in surface-proximal GLT-1 during cued reinstatement was not associated with dendrites from D1- (**L**, Kruskal-Wallis=0.7498 p=0.6874), or D2-MSNs (**M**, Kruskal-Wallis=25.49 p<0.0001, *p<0.05 vs. Sal). Like astrocyte-synaptic adjacency, GLT-1 co-registration with D2-dendrites was higher at baseline compared with D1-dendrites (p=0.0043 using Dunn’s test). In (**I**-**J**, **L-M**), N=25-44/4-5 cells/animals per group and full data spread is reported in Table S2. Sal, yoked saline; Ext, extinguished; Rst, 15-min cued reinstatement.

In contrast with the morphological adaptations, when data from D1- and D2-Cre rats were pooled the surface-proximal GLT-1 that co-registered with NAcore dendrites did not fully parallel the findings in Sprague-Dawley rats (Fig. 2C). Although GLT-1 was reduced in extinguished rats, the increase in surface-proximal GLT-1 after 15-min cued heroin seeking (Fig. 2D) was replicated in the pooled D1- and D2-Cre data (Fig. 6K). When dendritic adjacency of GLT-1 was examined separately in D1- and D2-Cre rats, levels of GLT-1 adjacent to D2 dendrites was higher when compared with D1 dendrites (compare saline-treated groups in Fig. 6L-M). The extinction-associated reduction in the pooled data occurred at D2-, not D1-dendrites (Fig 6L-M). However, the increase in surface-proximal GLT-1 after 15 min of cue exposure did not co-register with either D1 or D2-dendrites (Fig. 6L-M). Both the morphological plasticity and GLT-1 dynamics in NAcore astroglia from D1- and D2-Cre animals occurred in a manner that was not sex-specific (Fig. S6).

Taken together, these data show that the cue-induced increase in type 1 astroglia during heroin seeking (Fig. 3D-E) can be linked to morphological plasticity of astroglia surrounding D2-dendrites (Fig. 6H-J), but that the cue-induced increase in type 2 astroglia exhibiting high levels of extrasynaptic GLT-1 (Fig. 3D-E) are not associated with either D1- or D2-dendrites (Fig. 6K-M). A summary schematic illustrating subcircuit-selective astrocyte adaptations after extinction of heroin seeking is shown in Fig. S7.

## DISCUSSION

Synaptic glutamate spillover from prefrontal cortical synapses in NAcore is produced by drug associated cues and is necessary for drug cues to induce synaptic adaptations that regulate the intensity of drug seeking. Astroglia possess the capacity to strongly regulate glutamate spillover from synapses by their morphological proximity to synapses (*28*) and their expression of GLT-1, especially in perisynaptic astroglial processes (*29–32*). We show that during cue-induced heroin seeking, astroglia employ both of these mechanisms to dampen the intensity of seeking. Surprisingly, cue-induced increases in synaptic proximity and surface GLT-1 expression occurred in separate astroglial subpopulations. Type 1 astroglia had greater proximity to synapses in NAcore, while type 2 astroglia showed increased surface expression of GLT-1. The functional relevance of cues transiently inducing these astroglial subpopulations was shown by selectively eliminating either type 1 or type 2 astroglial plasticity and potentiating cue-induced heroin seeking. Finally, we demonstrated that the type 1 adaptation was selective for D2-, not D1-MSNs during seeking, while the cue-induced type 2 adaptation was not significantly associated with either D1- or D2-MSNs. Together, our data reveal at least two mechanisms whereby astroglia negatively regulate cue-induced motivation to seek heroin, but not sucrose. At least part of this action may arise from G_q_-coupled stimulation of astroglial processes toward D2-MSN synapses.

### Distinct subpopulations of astroglia during cue-induced heroin seeking

During cue-induced heroin seeking three distinct subpopulations of astroglia were present, type 1 showing increased synaptic proximity, type 2 showing upregulated surface GLT-1 and type 3 having relatively lower levels of both synaptic adjacency and surface GLT-1. Unbiased PCA revealed that type 1 and type 2 adaptations were in distinct subpopulations of astroglia. The appearance of type 1 and type 2 subpopulations induced by cued seeking was transient and absent after 120 min of unrewarded seeking. This is similar to the transient potentiation of D1-MSNs triggered by drug-associated cues (*26, 33*).

Control saline or sucrose rats showed largely type 1 and 3 astroglia. Type 2 astroglia emerged while type 1 were eliminated in heroin extinguished rats. Cue-induced heroin seeking was accompanied by transient return of type 1 astroglia to levels present in control rats and a transient elevation in the proportion of astroglia expressing the type 2 adaptation. In sum, our data clearly show that the appearance and disappearance of different types of astroglial adaptations depends on their stage in the addiction protocol (i.e. extinguished vs. relapsing) and reveals remarkable morphological and signaling plasticity in astroglia to negatively regulate cue-induced heroin seeking.

### Type 1 astroglia: Cue-induced morphological plasticity with no increase in surface GLT-1

A portion of mature synapses throughout the brain are insulated by astroglia (*34*), and the perisynaptic astroglial membrane expresses the highest density of GLT-1, glutamate receptors and other proteins, including actin binding proteins like ezrin, which together contribute to modulating glutamatergic synaptic activity. Perisynaptic astroglia modulate synaptic glutamate homeostasis in part through dynamic morphological plasticity in response to synaptic neurotransmission (*29, 34, 35*). In the hippocampus and cerebellum, the astroglial sheath is biased toward the postsynapse (*36*), permitting access of synaptically released glutamate onto presynaptic mGluR2/3 autoreceptors that negatively regulate release probability (*37*). Notably, dense expression of GLT-1 proximal to the synapse may prevent recruitment of this autoinhibitory mechanism, by retrieving glutamate from the synapse and thus reducing glutamate concentration at presynaptic autoreceptors. In this way, the GLT-1-deficient astroglial processes present in the type 1 subpopulation can promote autoinhibitory regulation of excitatory transmission in the NAcore by sterically guiding synaptically released glutamate towards presynaptic mGluR2/3 in the absence of glutamate uptake. The importance of guiding glutamate to presynaptic mGluR2/3 to negatively regulate relapse is supported by the fact that stimulating mGluR2/3 in the accumbens inhibits drug seeking (*38, 39*).

Astrocyte fine process motility is increased by activating G_q_-coupled mGluR5 (*15, 16*). Also, stimulating G_q_-DREADD in NAcore astroglia reduces reinstated cocaine seeking by promoting astroglial glutamate release onto presynaptic mGluR2/3 (*40*). Our findings complement this work by showing that activating astroglial G_q_-DREADD in the NAcore increases the number of synaptic puncta with near-adjacent astroglial processes, although the resolution limit of confocal microscopy does not permit us to determine whether astroglial processes were oriented toward the pre- or postsynapse after DREADD stimulation. Increasing G_q_ signaling in NAcore astroglia did not impact surface-proximal GLT-1 levels, supporting discrete signaling mechanisms for heroin cue-induced formation of type 1 and type 2 astroglial plasticity.

In addition to potentially guiding synaptically released glutamate toward mGluR2/3 autoreceptors, the synaptic re-association induced by heroin cues in type 1 astroglia may limit access of synaptic glutamate (i.e. glutamate spillover) to perisynaptic NMDA glutamate receptors containing the 2B subunit (NR2B) that must be stimulated for cues to initiate heroin seeking, and for the synaptic potentiation that accompanies heroin seeking (*33*). Finally, not only would synaptic insulation by astroglial processes be more likely to engage presynaptic autoinhibition and block access to NR2B necessary for synaptic potentiation, but synaptic adjacency of astroglial processes reduces the possibility of synaptic recruitment that occurs via transmitter spillover to neighboring synapses. This is supported by the fact that synaptic retraction of astroglial processes after LTP induction promotes synaptic crosstalk (*41*).

### Type 2 astroglia: Cue-induced increases in surface-proximal GLT-1 with no change in synaptic proximity

A portion of astroglia responded to heroin use and extinction by increasing surface-proximal GLT-1 and this adaptation was transiently accentuated by heroin cues. However, the increase in GLT-1 did not co-register with Synapsin I in either treatment group, indicating that surface GLT-1 in extinguished and reinstated animals was distant (>250 nm) from the synapse (i.e. extrasynaptic). Moreover, the increase in surface GLT-1 during seeking was not associated with either D1- or D2-dendrites. We hypothesize that type 2 astroglia do not guide glutamate spillover to presynaptic autoreceptors or sterically shield extrasynaptic high affinity glutamate receptors from synaptic glutamate, as discussed above for type 1 astroglia. However, diffusion of glutamate to more distal sites would be reduced by increased extrasynaptic GLT-1 (*22*). Thus, the primary impact of type 2 astroglia may be to deny access of synaptic glutamate to neighboring synapses thereby preventing synaptic recruitment. Notably, by inhibiting glutamate access to neuronal nitric oxide synthase (nNOS) interneurons where mGluR5 stimulation promotes NO-induced synaptic potentiation in D1-MSNs (*42*) or reducing NR2B stimulation on adjacent synapses (*33*), extrasynaptic GLT-1 could dampen cued heroin seeking and the associated synaptic potentiation (*43*).

While some studies demonstrate that the bulk of GLT-1 is on the astroglial surface (*31*), there are notable differences between hippocampal or cultured astrocytes and striatal astrocytes studied here (*44*). Our data indicate that a proportion of NAcore astroglia express relatively high levels of surface-proximal GLT-1 and that these cells target their GLT-1 extrasynaptically during relapse. Heterogeneity in GLT-1 expression by NAcore astroglia is depicted in Fig. S5 and is further supported by the non-Gaussian distribution of GLT-1 expression quantified on astrocytes from yoked saline control animals (Fig. 2B-C). Notably, type 2 astrocytes that exhibit high levels of surface-GLT-1, but low synaptic proximity were not abundant in control saline or sucrose groups, but emerged after extinction from heroin self-administration. Whether this adaptation was a consequence of heroin, heroin withdrawal or extinction training during withdrawal is a remaining question. However, it is noteworthy that extinction from sucrose self-administration did not increase the presence of type 2 astroglia, indicating that extinction train per se is not sufficient to produce this adaptation.

That G_q_ signaling did not stimulate plasticity in type 2 astroglia raises the possibility that levels and/or patterns of G_q_-coupled receptor expression may distinguish type 1 from type 2 astroglia. In keeping with this hypothesis, astroglial mGluR5 signaling stimulates peripheral process motility (*15*) and down-regulates GLT-1 (*45*). Such a dual role for mGluR5 signaling would contribute to the relatively low levels of surface GLT-1 in type 1 astroglia. Alternatively, stimulation of astroglial mGluR3 has been linked to increased GLT-1 expression (*45*), raising the possibility that cue-induced glutamate spillover may act via receptors differentially expressed on type 1 and 2 astroglia.

### Astroglial adaptations are differentially regulated around D1- and D2-MSNs

D1- and D2-MSNs together constitute 90-95% of all NAcore neurons (*9*) and different astroglial subpopulations modulate synaptic transmission selectively on one or the other neuronal subtype (*19, 27*). Consistent with astroglia associating selectively with one or the other, but not both types of MSN, astroglia were more proximal to D2-than D1-MSN dendrites in control animals. The finding that astroglia increased their proximity to D1-MSN synapses after extinction training, but retracted from D2-MSNs, is consistent with the fact that synaptic adaptations associated with extinction from addictive drug self-administration are found primarily on D2-MSNs, while adaptations produced by drug cues are predominant on D1-MSNs (*26, 46–48*). We predict that astrocyte insulation of D1-MSNs after extinction training would reduce synaptic glutamate spillover, extrasynaptic receptor simulation and ultimately synaptic potentiation of D1-synapses during extinction, relative to D2-MSNs (*26, 48*). Moreover since astrocytes can trigger long-term depression at D1-MSN synapses via mGluR5-dependent ATP/adenosine signaling in the dorsal striatum (*19*), an increase in astrocyte association with D1-MSN synapses after heroin extinction may serve to suppress synaptic activity at D1-neurons and to thereby dampen seeking. Conversely, retraction of astrocytes from D2-MSNs after extinction training is expected to be permissive of synaptic potentiation at D2-MSN synapses via glutamate spillover. A number of studies are consistent with this interpretation, including: extinction from cocaine is associated with synaptic potentiation at D2- but not D1-MSN excitatory synapses (*26*), matrix metalloprotease-2 (MMP-2) activity is elevated in heroin extinguished rats only around NAcore D2-MSN dendritic spines (*46*), and chemogenetically activating D2-MSNs suppresses drug seeking (*10*).

That astrocyte processes surrounding D1-MSNs retracted during heroin-associated cue exposure is consistent with permitting glutamate spillover and potentiating synaptic activity at D1-synapses, as has been reported for both glutamate transmission and spine morphology (i.e. increased spine head diameter and/or spine density) during cued drug seeking (*26, 47*). Since astrocytes dampen synaptic potentiation via a number of mechanisms (*20, 40*), we hypothesize that increased astrocyte insulation of D2-MSNs during cue-reinstated heroin seeking may reduce D2-MSN potentiation relative to the extinguished context (*26, 41*). However, a number of reports indicate that subpopulations of D2-MSNs in the striatum promote, rather than inhibit motivated behaviors (*49–52*). Moreover, cortical inputs linked to cued reinstatement (*53*) innervate D2-MSNs to a greater extent than D1-MSNs (*54*). For this reason, the possibility remains that increased astrocyte insulation on D2-MSNs during cued reinstatement may serve a compensatory function by triggering autoinhibition at cortical terminals that release glutamate in response to heroin-conditioned cues. In this way, astrocyte insertion onto D2-MSNs may contribute to reducing glutamate spillover onto both D2-MSNs, as well as D1-MSNs, that receive less direct innervation by prefrontal terminals (*54*), but are significantly engaged during cued reinstatement to drive seeking (*26*).

Akin to astroglial synaptic proximity, surface GLT-1 was lower adjacent to D1-compared with D2-dendrites in saline rats. However, the overall dynamic in type 2 astroglia after extinction and cued reinstatement observed in wild-type rats was not recapitulated in either D1- or D2-MSNs. This could result from increased surface-GLT-1 in type 2 astroglia targeting non-dendritic portions of D1- and D2-MSNs or targeting subpopulations of interneurons or other astroglia that were not labeled in D1- and D2-Cre rats.

### Astroglial plasticity during sucrose versus heroin seeking

Sucrose training did not down-regulate GLT-1 nor did it produce changes in synaptic proximity by NAcore astroglia in any treatment group (*6, 55*). The relative lack of changes in astroglial subpopulation plasticity after extinction from sucrose self-administration and cued reinstatement supports the likelihood that changes in the proportion of astroglial subtypes in heroin trained rats may be selective for addictive drugs and not natural rewards. Interestingly, although we found no change in co-registration of GLT-1 with Synapsin I in sucrose-trained or reinstated animals, we observed reduced surface expression of GLT-1 after extinction from sucrose self-administration compared to yoked controls, further reducing the proportion of type 2 astroglia in this group. As described above, we predict that synaptic adjacency of astroglial processes deficient in GLT-1 would favor presynaptic autoinhibition upon transmitter release (*36*). Consistent with this hypothesis, sucrose-trained rats exhibit more rapid extinction training and less perseverative reward seeking compared with drug-trained animals (*56, 57*), even though during sucrose self-administration produced more active lever presses than heroin self-administration.

### Implications for treating substance use disorders

We found that astroglia in the NAcore exhibited two forms of plasticity capable of shaping cue-induced heroin seeking that occurred in separate astroglial subpopulations and were regulated by distinct signaling mechanisms. Moreover, both increasing astroglial proximity to synapses and glutamate transport involve proteins selectively expressed in astroglia (ezrin and GLT-1, respectively). The fact that none of the plasticity produced in NAcore astroglia from heroin-trained rats was recapitulated by sucrose training supports the potential that selective interventions in one or the other astroglial dynamic may be therapeutically beneficial in treating substance use disorder. Indeed, pharmacological treatments such as ceftriaxone and N-acetylcysteine that elevate GLT-1 are effective at reducing drug seeking in rodent models of relapse (*58–60*) and reducing drug cue reactivity in humans (*61*). Unfortunately, while reducing craving induced by drug cues, N-acetylcysteine has proven only marginally effective at reducing relapse in human trials (*62–64*). Given that astroglia engage two distinct processes for dampening cue-induced drug seeking, and that increasing synaptic adjacency of astroglial processes in the absence of GLT-1 reduces the intensity of relapse, the poor efficacy in relapse prevention by drugs restoring GLT-1 might be improved in combination with drugs that promote synaptic adjacency of NAcore astroglia.

## MATERIALS AND METHODS

### Experimental design and statistical analyses

All experiments included groups with ≥ 4 animals each to minimize variability due to animal behavior. All immunohistochemical and imaging procedures using tissue from experimental groups (i.e. extinguished, reinstated) were conducted alongside yoked controls to minimize the impact of experimental variability on independent groups. Data were analyzed using GraphPad Prism and a D’Agostino-Pearson normality test followed by Kruskal-Wallis or Mann-Whitney tests when one or more groups were not normally distributed. For non-Gaussian datasets, including all astroglial measures, no outliers were removed and data were plotted with values for each cell depicted individually and a bar to indicate the group median. Dunn’s test was used for post hoc comparisons. Normally distributed datasets, including all behavioral data, were analyzed using a 1- or 2-way ANOVA or a Student’s t-test. To more clearly show median differences in Fig. 6, y-axes show data ≤2.0% co-registration. Full datasets for Fig. 6 are reported in Table S1. Cumulative distributions were analyzed using Kolmogorov-Smirnov or Chi^2^ and a Bonferroni correction was applied for multiple comparisons. In all cases, *p* <0.05 was considered significant.

### Self-administration

Experimental procedures involving animals were conducted in accordance with guidelines established by the Institutional Animal Care and Use Committee at the Medical University of South Carolina. Operant training was conducted as previously described (*6*). Briefly, male Sprague Dawley rats or male and female Long Evans rats (200-250g) were anesthetized with i.m. ketamine (100 mg/kg) and xylazine (7 mg/kg) and fitted with intrajugular catheters. Rats were trained to self-administer heroin during 3h sessions for 10d and presses on an active lever were paired with light and tone cues and i.v. heroin infusion. Animals trained to self-administer sucrose did not undergo catheter implantation and received sucrose (45 mg, Bio-Serv) in place of heroin along with cues during self-administration (2h/d). Yoked controls were played cues when a paired rat received heroin or sucrose. Rats yoked to heroin self-administering animals also received i.v. saline infusions. After self-administration, animals underwent 10-12d of extinction training (3h/d after heroin, 2h/d after sucrose) where active lever presses yielded no reward or cues. Extinguished rats and yoked controls were sacrificed 24h after the final extinction session. Reinstated animals were placed in the operant chamber for 15 or 120m and cues were restored to the active lever, but no reward was delivered.

### Viral labeling

After catheter implantation or 5d before starting sucrose self-administration, rats received microinjections (1 μL/hemisphere, 0.15 μL/minute, 5 min diffusion) of a virus driving expression of membrane-targeted mCherry under control of the GFAP promoter (AAV5/GFAP-hM3dq-mCherry, University of Zurich, Figs. 1-5 or AAV5/GfaABC1D-Lck-GFP, Addgene, Fig. 6) in the NAcore (+1.5mm AP, ±1.8mm ML, -7.0mm DV). To label D1- and D2-MSN dendrites, AAV1/CAG-Flex-Ruby2sm-Flag (Addgene) was co-injected in D1- or D2-Cre rats along with virus used to label astroglia. Virus incubation occurred over the course of operant training.

### Confocal imaging and image analysis

Animals were anesthetized with an overdose of pentobarbital (20 mg i.v. or 100 mg i.p.) and perfused transcardially with 4% PFA. Brains were incubated overnight in 4% PFA and sliced at 100 μm using a vibratome (Thermo Fisher). Slices containing the NAcore were permeabilized in PBS with 2% Triton X-100 for 1h at room temperature. Non-specific epitope binding was blocked by incubation in PBS with 0.2% Triton X-100 (PBST) and 2% NGS (block) for 1h at room temperature before incubation in primary antibodies (rabbit anti-Synapsin I, ab64581, Abcam; guinea pig anti-GLT-1, ab1783, EMD Millipore; mouse anti-FLAG, F1804, Sigma-Aldrich) at 1:1000 in block for 48h. After washing in PBST, tissue was incubated overnight in biotinylated anti-guinea pig antibody (BA-7000, Vector Laboratories) in PBST and then overnight in fluorescently-labeled antibodies (Thermo Fisher) in PBST after washing. Tissue was mounted onto glass slides before imaging with a Leica SP5 laser scanning confocal microscope. All images were acquired at 63x using an oil immersion objective lens, 1024 x 1024 frame size, 12-bit resolution, 4-frame averaging and a 1-μm step size. Z-stacks were iteratively deconvolved 10 times (Autoquant) and digital analysis of astroglial mCherry or GFP signal intensity relative to background was used to generate a digital model of each astrocyte (Bitplane Imaris). Rendered astrocytes were used to mask Synapsin I and GLT-1 or Flag signal that was not co-registered with the astroglial volume. Co-registration (astrocyte with Synapsin I, astrocyte with GLT-1, astrocyte with GLT-1 and Synapsin I, astrocyte with Flag and Synapsin I, astrocyte with Flag and GLT-1) was determined based on thresholded signal intensity in each channel. Voxels containing signal intensity greater than noise in each channel were determined empirically using the colocalization module and were used to build a colocalization channel. The surface module was then used to determine the net volume of co-registered signal. All measures, with the exception of surface-proximal GLT-1 (Fig. 2C-D, Fig. S1C-D), and synaptic and extrasynaptic GLT-1 (Fig. 2F-G, Fig. SF-G), were normalized to the volume of the astrocyte from which they were generated to control for changes in astroglial volume (*6*) and to permit inclusion of astrocytes that were cropped along the z-axis during imaging. Surface-proximal GLT-1 was determined by excluding co-registered signal that was within the astrocyte volume, but >250nm (a distance roughly equivalent to the lateral resolution of the microscope used for imaging) from the membrane and was normalized to total GLT-1 from the astroglial volume. Synaptic and extrasynaptic GLT-1 values were normalized to surface GLT-1 levels from each astrocyte. In all case, imaging and analyses were conducted blind to animal treatment.

### Principal component analysis and hierarchical clustering

To identify astroglial clusters according to synaptic co-registration as percent of astrocyte volume, and surface-proximal GLT-1 expression as percent of total GLT-1 expression, hierarchical clustering on principal components was computed using R software (R Core Team, 2020) according to (*21*). Principal components were computed along the two specified dimensions and Ward’s criterion was applied on selected principal components (Fig. S3). The output dendogram was separated along 2 main breaks in the data to form 3 clusters (Fig. S3).

### G_q_-DREADD stimulation

AAV5/GFAP-hM3dq-mCherry (University of Zurich) was diluted 1:10 in sterile saline and delivered to the NAcore (+1.5mm AP, ±1.8mm ML, -7.0mm DV) while rats were under anesthesia for jugular catheter placement as in (*40*). This dilution was used, because it produced sufficient astroglial labeling with little off-target labeling in neurons (Fig. 4A). Viral incubation occurred over the course of operant training. Twenty-four hours after the last extinction session, rats received CNO (3mg/kg i.p., Abcam) or vehicle (5% DMSO in sterile saline) 30 minutes prior to perfusion and brain extraction.

### Vivo-morpholino knockdown

After catheter placement, animals were fitted with bilateral cannulae above the NAcore (+1.5mm AP, ±1.8mm ML, -5.5mm DV). Starting on day 6 of extinction training, animals received infusions of an ezrin antisense vivo-morpholino oligo (*6*), a GLT-1 antisense vivo-morpholino oligo (*24*), or a control oligo (Gene Tools, Inc.) 1.5mm beyond the base of the guide cannulae (50μmol/L, 1μL/hemisphere, 0.5μL/min) for 3 consecutive days. After 3 additional days of extinction training, rats underwent cued reinstatement for 15-min prior to sacrifice.

## Author contributions

A.K. and P.W.K. designed and interpreted the study and wrote the paper. A.K. conducted experiments with assistance from A.A. and C.G-K. H.L. performed PCA on astrocyte data generated by A.K.

## Data and materials availability

All data are available within the main text or the supplementary material.

---

**Fig. S1.**
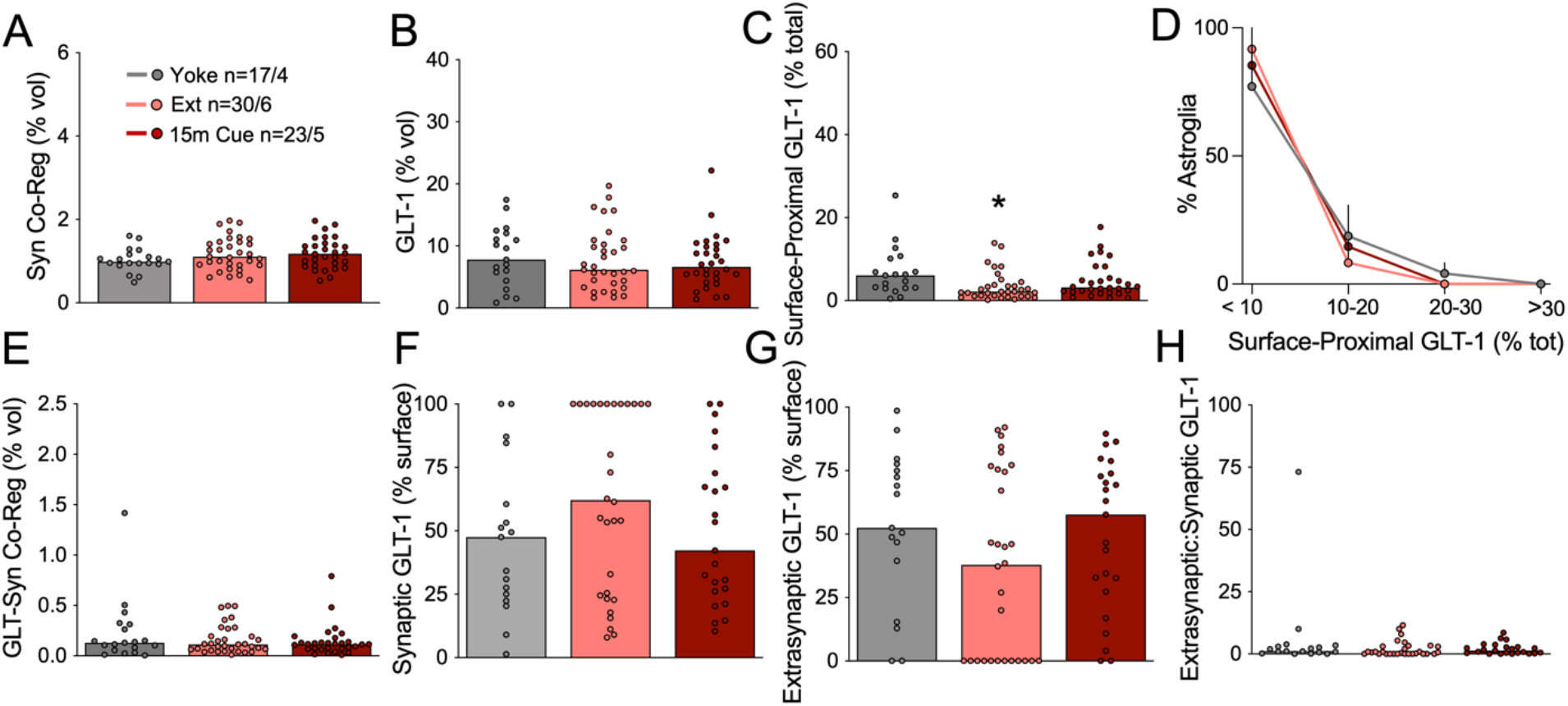
Astrocyte motility and GLT-1 expression were unchanged during reinstated sucrose seeking. Co-registration of labeled NAcore astroglia with Synapsin I was not changed after extinction of sucrose self-administration or during 15-min of cue-reinstated sucrose seeking (**A**, Kruskal-Wallis=2.153, p=0.341). (**B**) GLT-1 expression was unchanged after operant training with sucrose (Kruskal-Wallis=0.7336, p=0.693). (**C**) Surface-proximal GLT-1, shown as percent of total GLT-1 from each astrocyte, was reduced after extinction from sucrose self-administration (Kruskal-Wallis=8.056, p<0.05). (**D**) Cued reinstatement of sucrose seeking did not change the proportion of astroglia expressing high levels of surface-proximal GLT-1 (2-way ANOVA, F_6,36_=0.469, p=0.827). (**E**) Co-registration of GLT-1 with the presynaptic marker Synapsin I was not changed by sucrose training (Kruskal-Wallis=0.3724, p=0.830). When synaptic (**F**, Kruskal-Wallis=2.950) and extrasynaptic (**G**, Kruskal-Wallis=2.950) fractions of surface GLT-1 were analyzed separately, we found no change in sucrose-trained rats compared with yoked controls. (**H**) shows ratio of extrasynaptic:synaptic GLT-1 (Kruskal-Wallis=2.950). N shown in (**A**) as cells/animals. *p<0.05 compared to yoked control using Dunn’s test. Yoked, yoked cues; Ext, extinguished; 15m Cue, 15-min cued reinstatement.

**Fig. S2.**
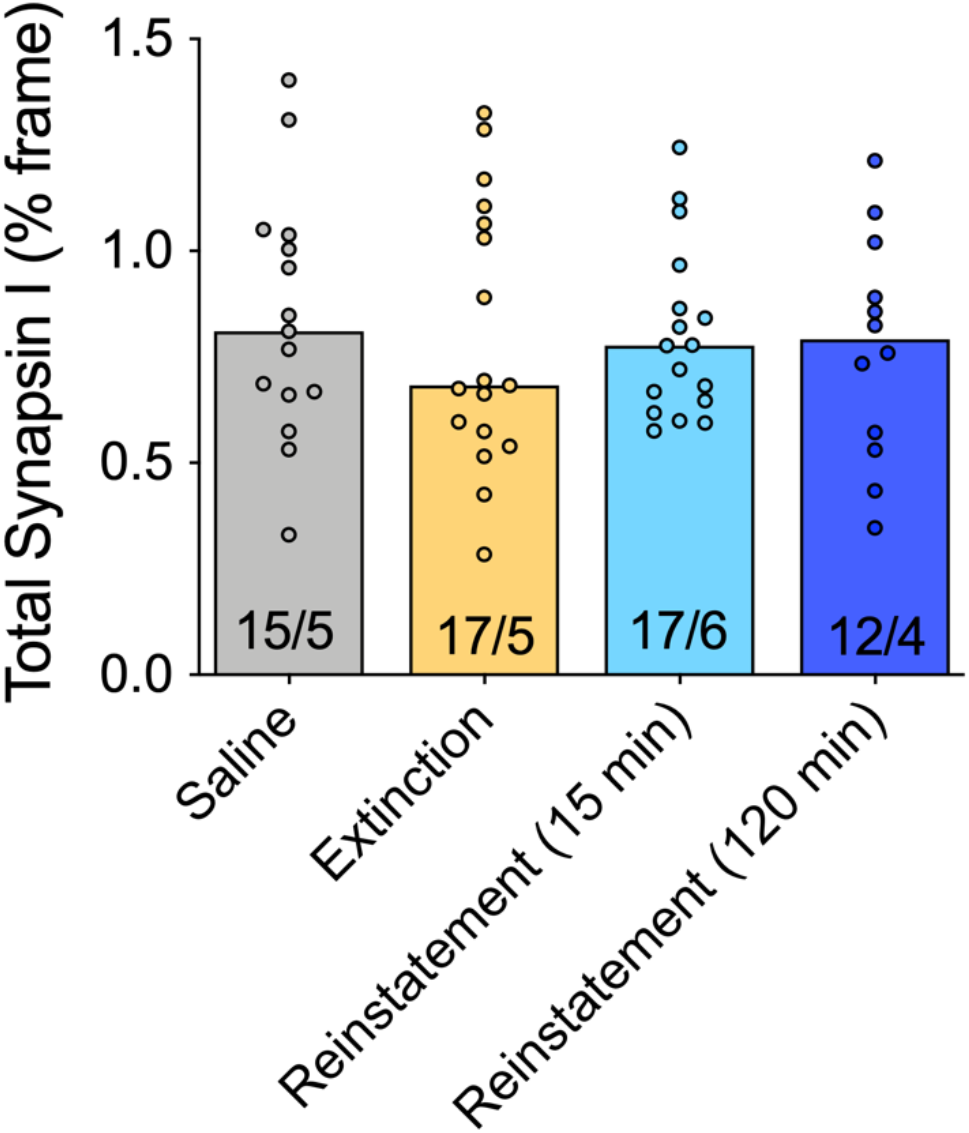
Heroin self-administration and extinction training did not impact Synapsin I immunoreactivity in the NAcore. Kruskal-Wallis=0.4046, p=0.9393. N shown in bars as frames/animal.

**Fig. S3.**
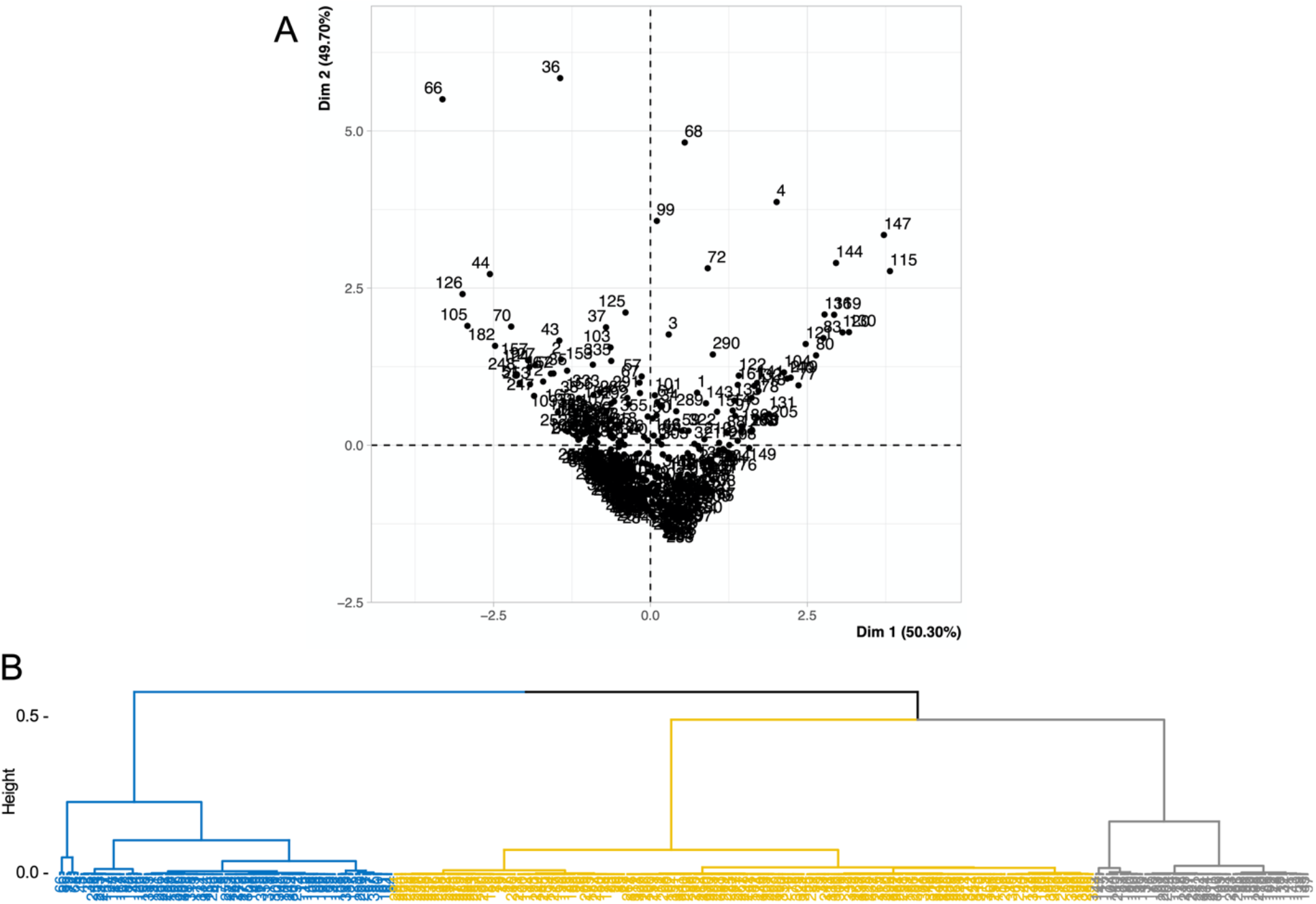
(**A**) Principal component analysis showing individual data points organized according to dimensions 1 and 2, which account for 50.3% and 49.7% of the data variance, respectively. (**B**) Dendrogram shows three clusters representing type 1 (blue), type 2 (gray), and type 3 (yellow) astroglia.

**Fig. S4.**
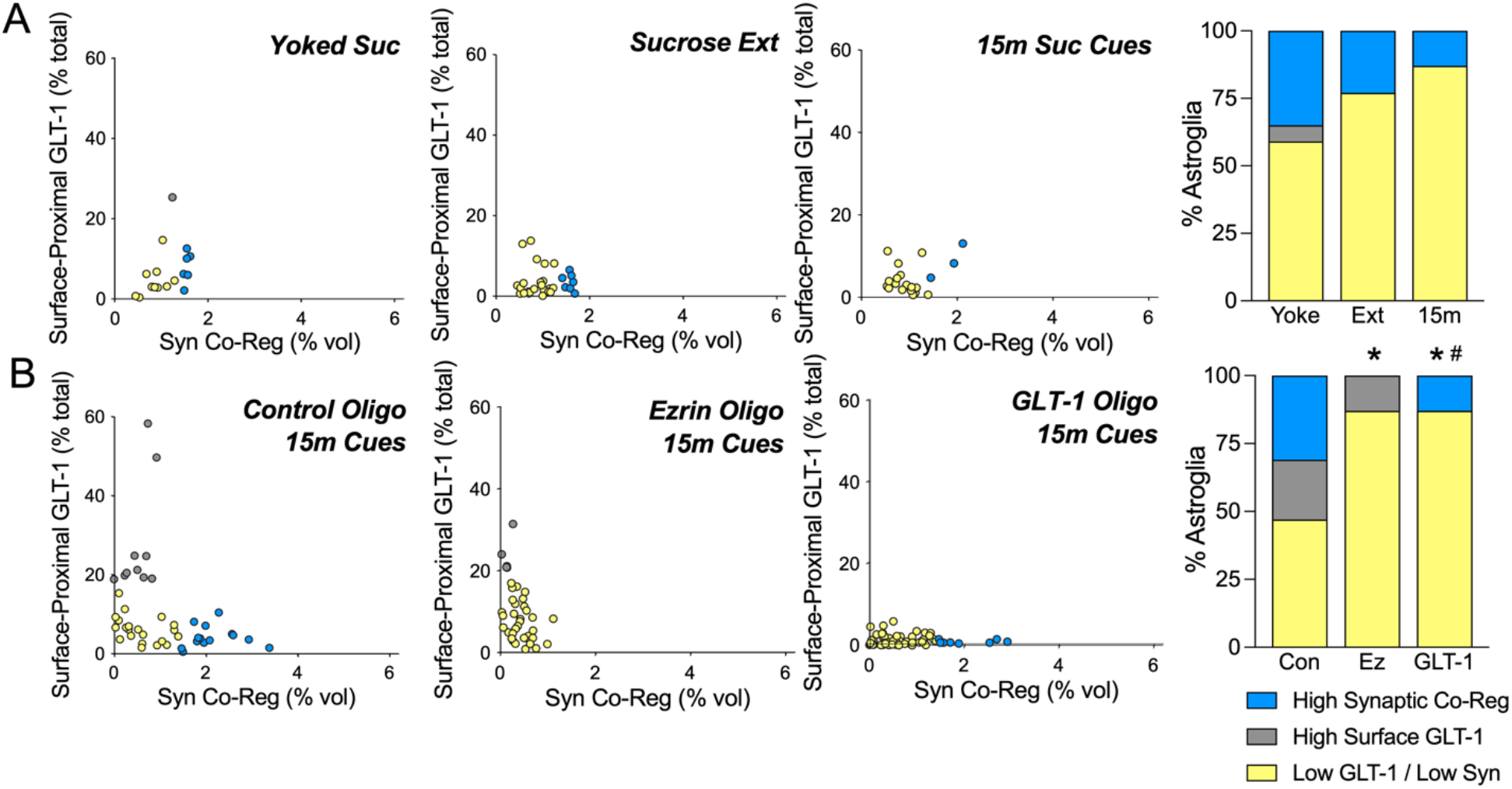
Astroglial subpopulations are not altered by operant training with sucrose, but are abolished by ezrin or GLT-1 oligo treatment. (**A**) Astroglial clusters are not altered by operant training with sucrose (Chi^2^=2.818 p=0.2444 Yoke vs. Ext, Chi^2^=4.535 p=0.1036 Yoke vs. 15m). (**B**) Compared with astroglial subpopulations during reinstatement after control oligo treatment (left), ezrin or GLT-1 oligo treatment abolished subpopulations characterized by high synaptic adjacency (blue) or high surface-proximal GLT-1 (gray), respectively (Chi^2^=17.87 *p=0.0002 Ez vs. Con, Chi^2^=23.60 *p<0.001 GLT-1 vs. Con, Chi^2^=13.01 #p=0.003 GLT-1 vs. Ez). In (**A**, right panel), Yoke, yoked cues; Ext, extinguished; 15m, 15-min cued reinstatement. In (**B**, right panel), Con, control oligo; Ez, ezrin oligo; GLT-1, GLT-1 oligo.

**Fig. S5.**
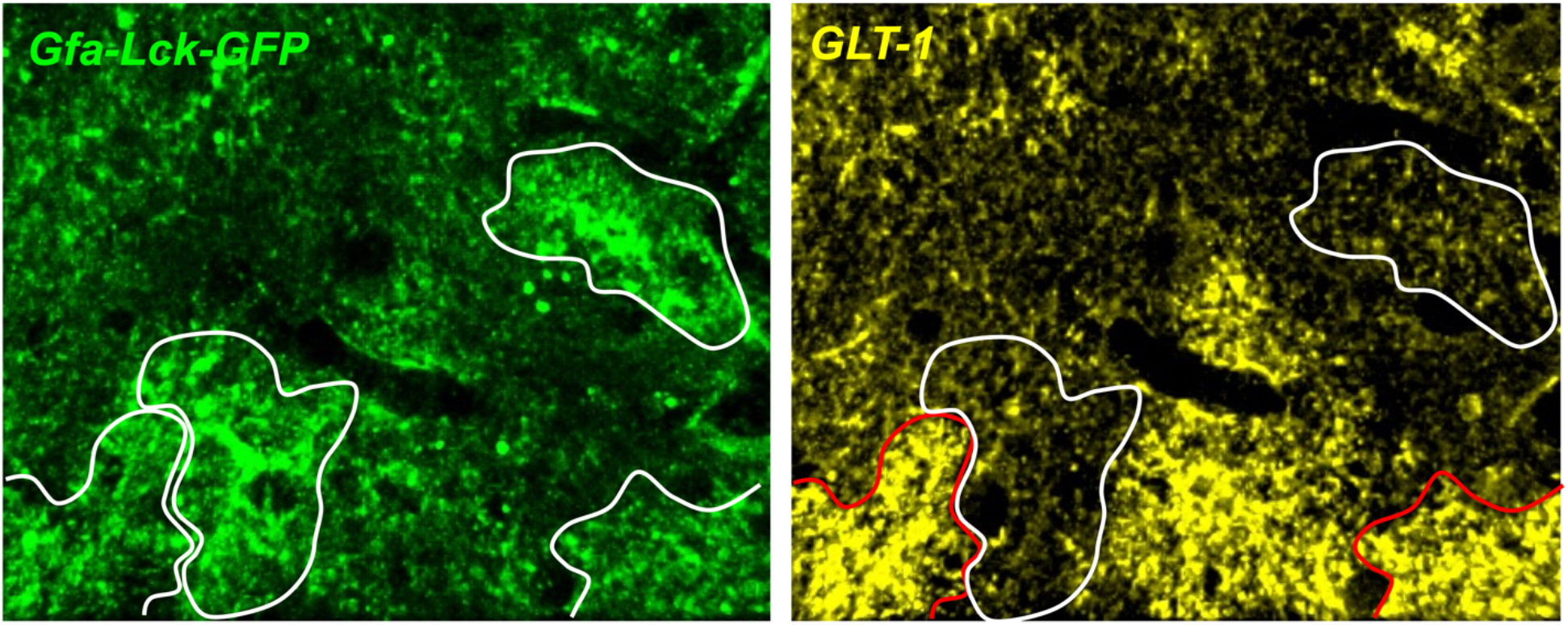
Astroglial transduction with AAV5/GfaABC1D-Lck-GFAP labels the astroglial membrane (green). Astrocytes identified using this marker are outlined in white (left panel). (Right panel) In the same frame, immunolabeling for GLT-1 (yellow) shows astrocytes with high (red outline) and low levels of GLT-1 expression (white outline).

**Fig. S6.**
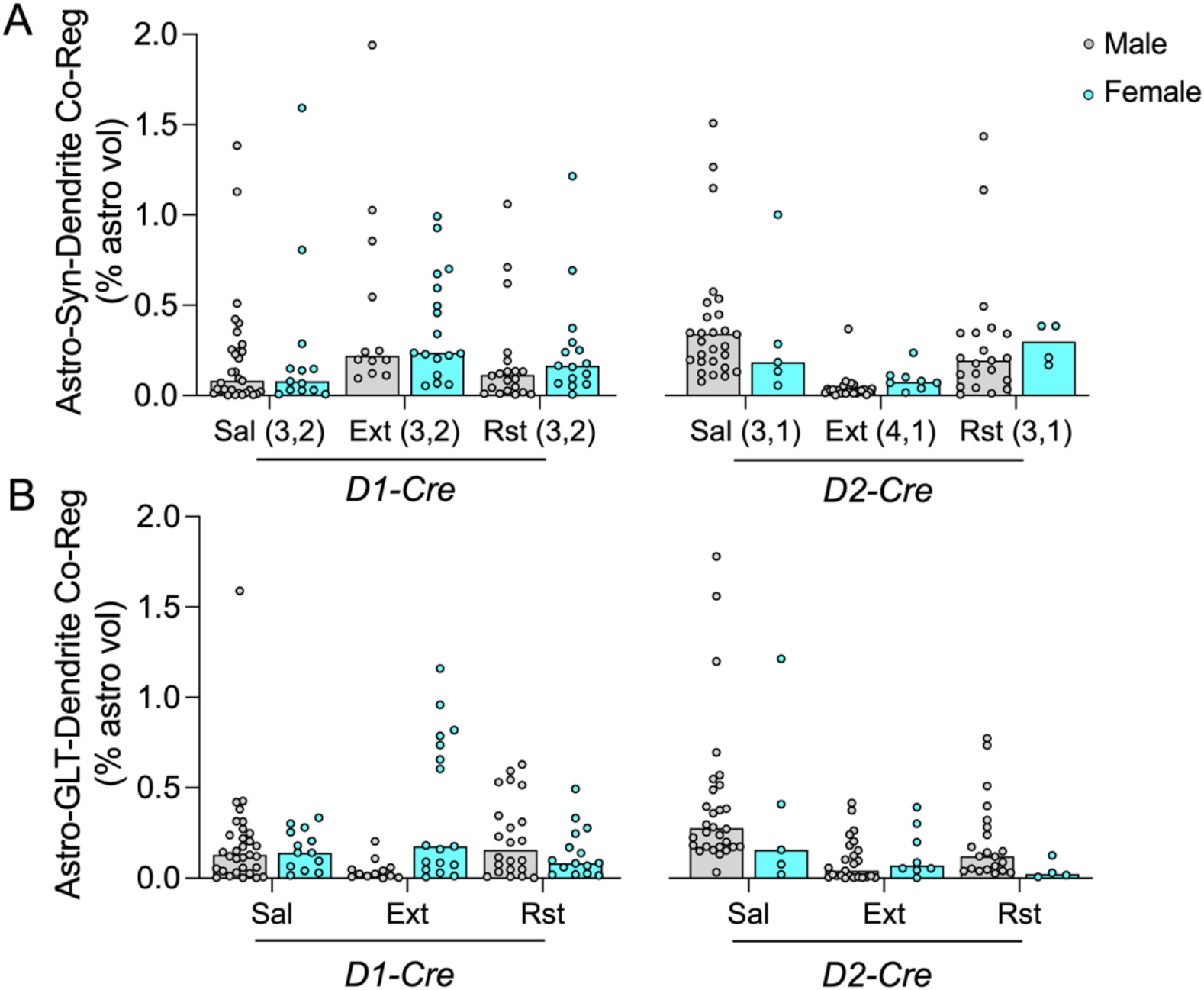
Cell-type specific dynamics in astrocyte-synapse and GLT-dendrite association are not sex-dependent. (**A**) No sex differences were detected in D1- or D2-synaptic co-registration by NAcore astroglia after operant training with heroin (2-way ANOVA F_1,196_=0.7529 p=0.3866). (**B**) Dendritic association of GLT-1 did not differ by sex (2-way ANOVA F_1,196_=0.05632 p=0.8127). Sal, yoked saline; Ext, extinguished; Rst, 15-min cued reinstatement. Animal N shown below bars in (**A**) as (male, female).

**Fig. S7.**
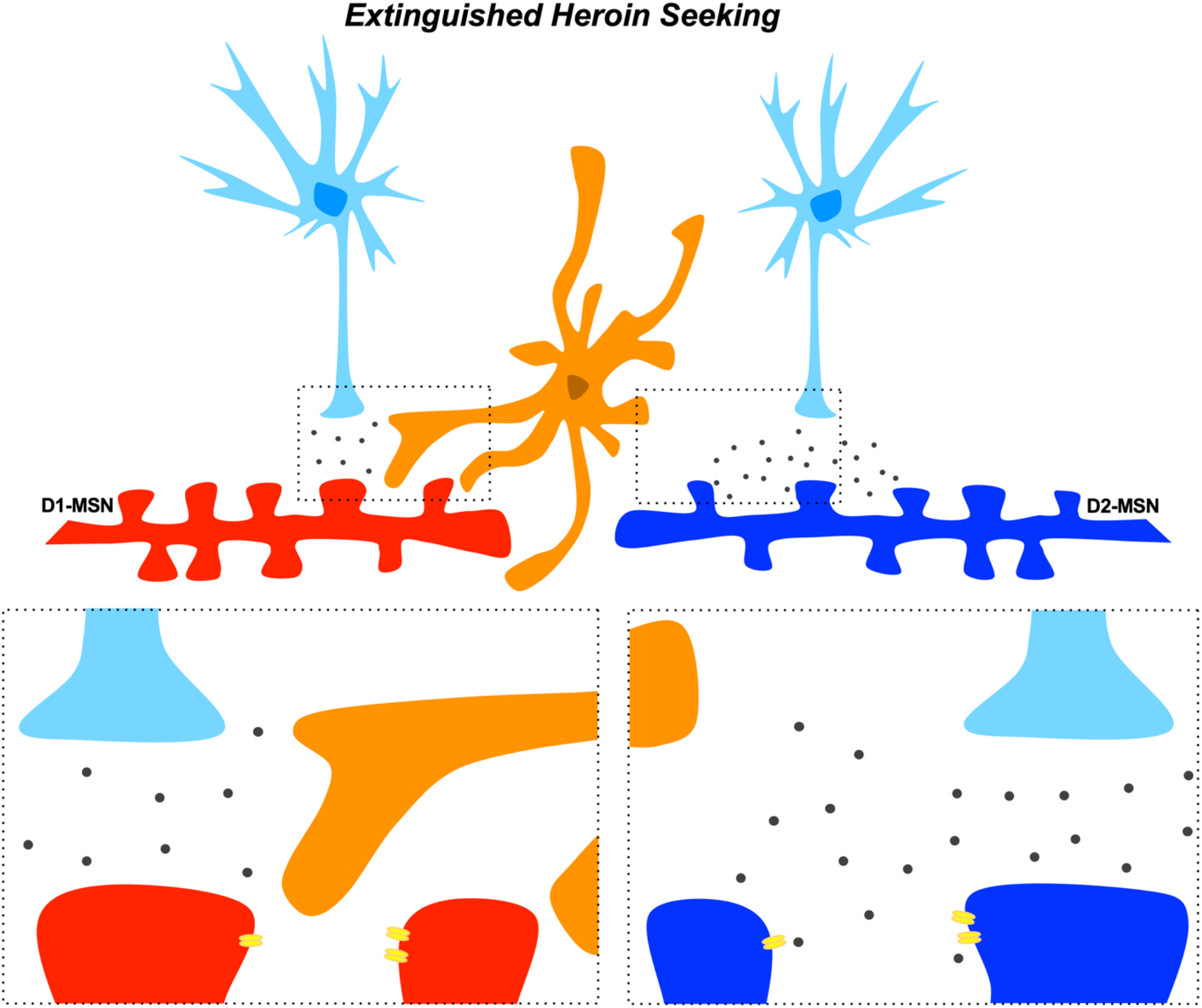
Synapse-selective proximity of astrocytes regulates seeking. Extinguished heroin seeking is characterized by astrocytes exhibiting a high degree of association with synapses from D1-MSNs (left), but retraction from synapses from D2-MSNs (right). High synaptic co-registration by astroglial processes low in GLT-1 (orange) is predicted to induce autoinhibition through spatial buffering of glutamate toward presynaptic inhibitory autoreceptors (*36*). The high co-registration of astroglia with D1-MSN synapses during extinction training is also predicted to shield post-synaptic NR2B receptors (yellow) that produce postsynaptic potentiation when stimulated by glutamate (*33*). These two functions would suppress D1-MSN potentiation during extinction training. Instead, retraction from D2-MSNs after extinction of heroin seeking engages post-synaptic potentiation of D2-MSNs through stimulation of postsynaptic NR2B (*33*), synaptic recruitment (*41*), and loss of autoinhibitory mechanisms at terminals synapsing onto D2-MSNs (*36*). These hypotheses are supported by data showing potentiation of D2-MSNs, but not D2-MSNs during extinction training (*26, 46*). Reversal of this pattern during cued reinstatement would permit potentiation of D1-MSNs and block signaling through D2-MSNs, permitting seeking behavior (*10*).

**Table S1.**
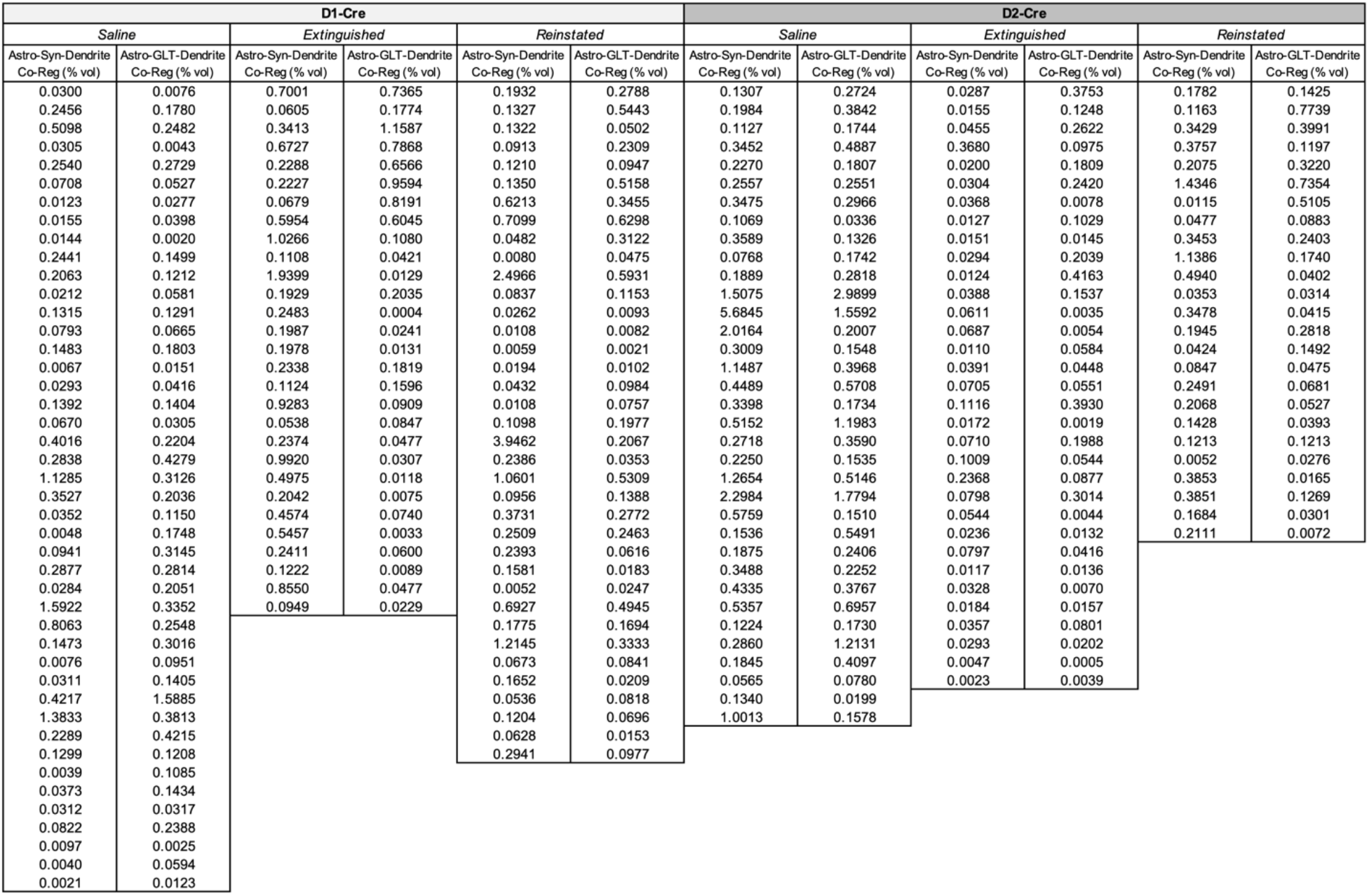
Dendrite-specific co-registration values. Triple co-registration of astroglia with Synapsin I and labeled dendrites, as well as triple co-registration of astroglia with GLT-1 and labeled dendrites from D1- and D2-Cre rats are presented as % astrocyte volume.

