## Supplementary Material for "Plasticity in astrocyte subpopulations regulates heroin relapse"

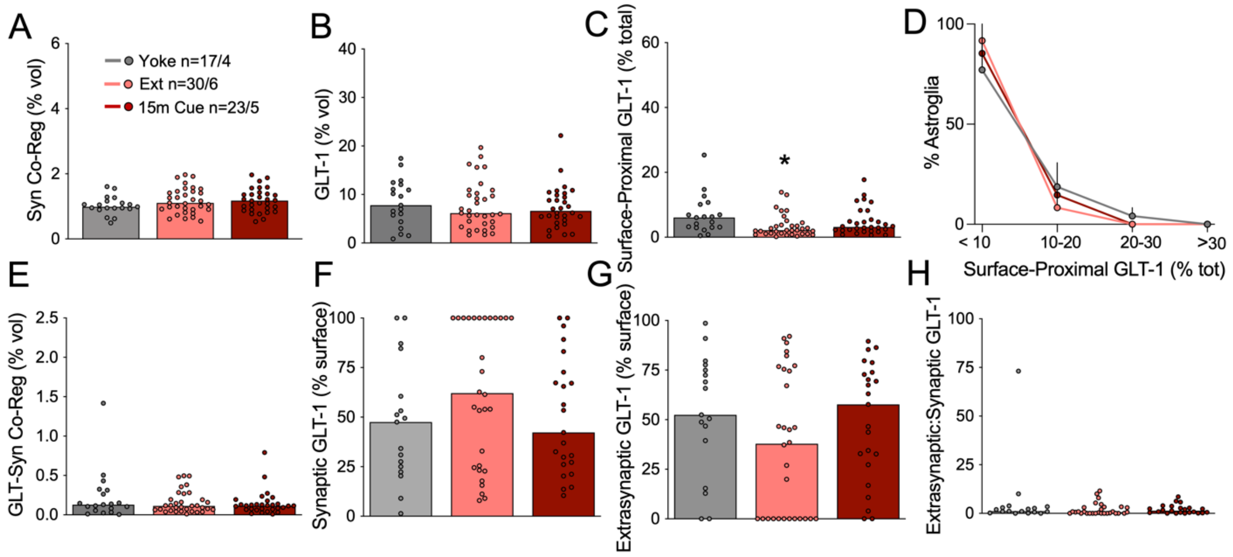


**Fig. S1. Astrocyte motility and GLT-1 expression were unchanged during reinstated sucrose seeking.** Co-registration of labeled NAcore astroglia with Synapsin I was not changed after extinction of sucrose self-administration or during 15-min of cue-reinstated sucrose seeking (**A**, Kruskal-Wallis=2.153, p=0.341). (**B**) GLT-1 expression was unchanged after operant training with sucrose (Kruskal-Wallis=0.7336, p=0.693). (**C**) Surface-proximal GLT-1, shown as percent of total GLT-1 from each astrocyte, was reduced after extinction from sucrose self-administration (Kruskal-Wallis=8.056, p<0.05). (**D**) Cued reinstatement of sucrose seeking did not change the proportion of astroglia expressing high levels of surface-proximal GLT-1 (2-way ANOVA, $F_{6,36}$=0.469, p=0.827). (**E**) Co-registration of GLT-1 with the presynaptic marker Synapsin I was not changed by sucrose training (Kruskal-Wallis=0.3724, p=0.830). When synaptic (**F**, Kruskal-Wallis=2.950) and extrasynaptic (**G**, Kruskal-Wallis=2.950) fractions of surface GLT-1 were analyzed separately, we found no change in sucrose-trained rats compared with yoked controls. (**H**) shows ratio of extrasynaptic:synaptic GLT-1 (Kruskal-Wallis=2.950). N shown in (**A**) as cells/animals. *p<0.05 compared to yoked control using Dunn's test. Yoked, yoked cues; Ext, extinguished; 15m Cue, 15-min cued reinstatement.

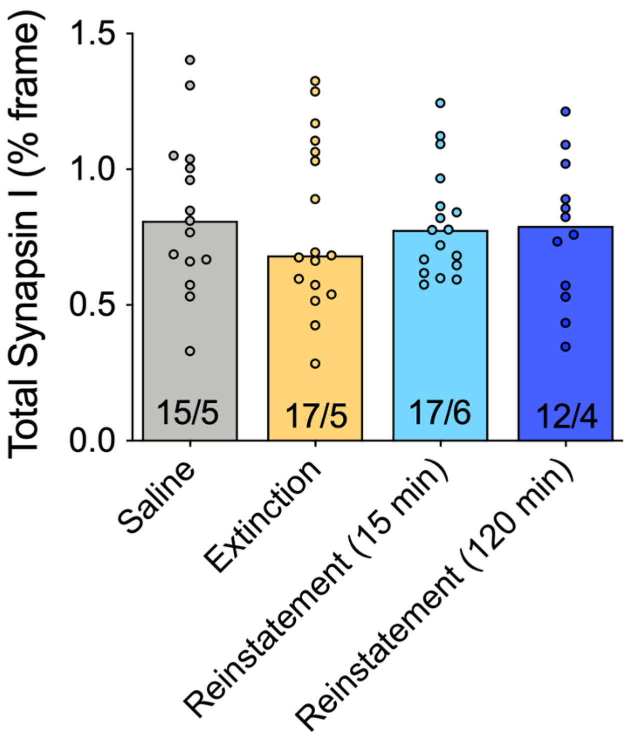


**Fig. S2. Heroin self-administration and extinction training did not impact Synapsin I immunoreactivity in the NAcore.** Kruskal-Wallis=0.4046, p=0.9393. N shown in bars as frames/animal.

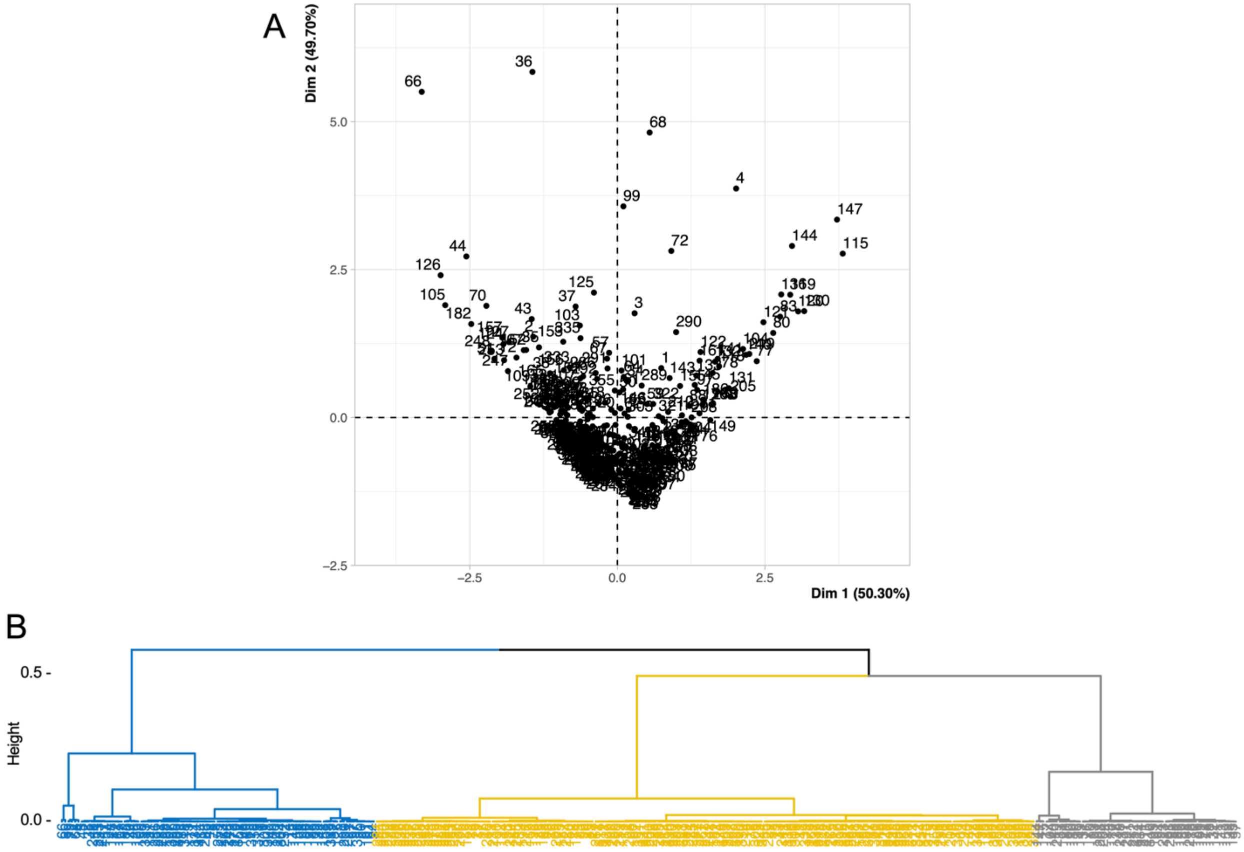


**Fig. S3.** (**A**) Principal component analysis showing individual data points organized according to dimensions 1 and 2, which account for 50.3% and 49.7% of the data variance, respectively. (**B**) Dendrogram shows three clusters representing type 1 (blue), type 2 (gray), and type 3 (yellow) astroglia.

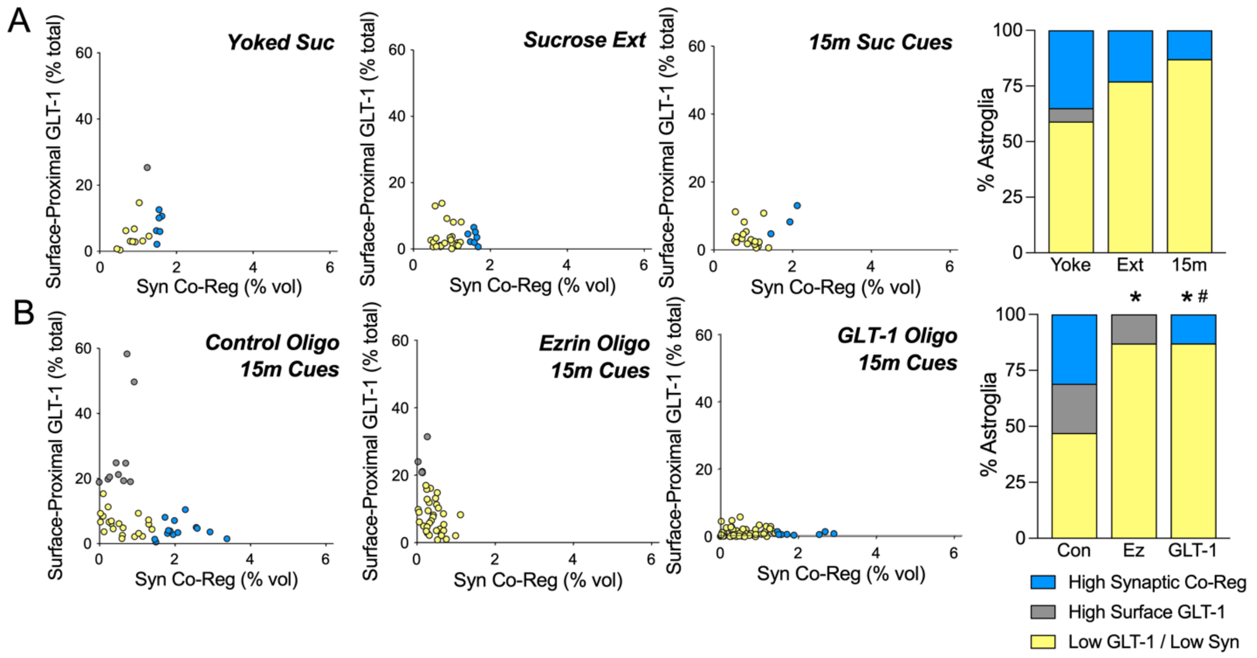


**Fig. S4. Astroglial subpopulations are not altered by operant training with sucrose, but are abolished by ezrin or GLT-1 oligo treatment.** (**A**) Astroglial clusters are not altered by operant training with sucrose (Chi$^2$=2.818 p=0.2444 Yoke vs. Ext, Chi$^2$=4.535 p=0.1036 Yoke vs. 15m). (**B**) Compared with astroglial subpopulations during reinstatement after control oligo treatment (left), ezrin or GLT-1 oligo treatment abolished subpopulations characterized by high synaptic adjacency (blue) or high surface-proximal GLT-1 (gray), respectively (Chi$^2$=17.87 *p=0.0002 Ez vs. Con, Chi$^2$=23.60 *p<0.001 GLT-1 vs. Con, Chi$^2$=13.01 #p=0.003 GLT-1 vs. Ez). In (**A**, right panel), Yoke, yoked cues; Ext, extinguished; 15m, 15-min cued reinstatement. In (**B**, right panel), Con, control oligo; Ez, ezrin oligo; GLT-1, GLT-1 oligo.

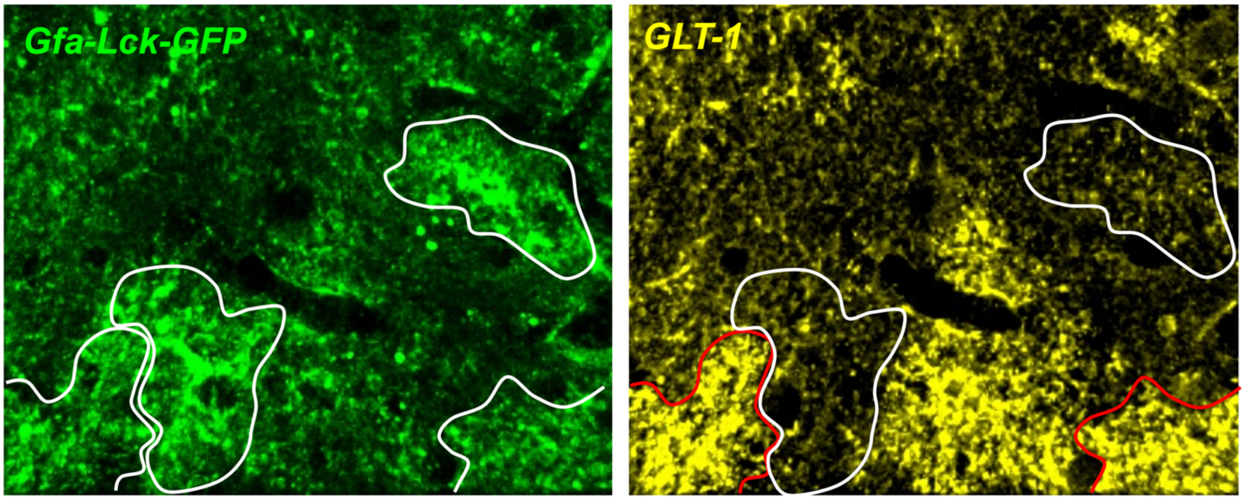


**Fig. S5.** Astroglial transduction with AAV5/GfaABC1D-Lck-GFAP labels the astroglial membrane (green). Astrocytes identified using this marker are outlined in white (left panel). (Right panel) In the same frame, immunolabeling for GLT-1 (yellow) shows astrocytes with high (red outline) and low levels of GLT-1 expression (white outline).

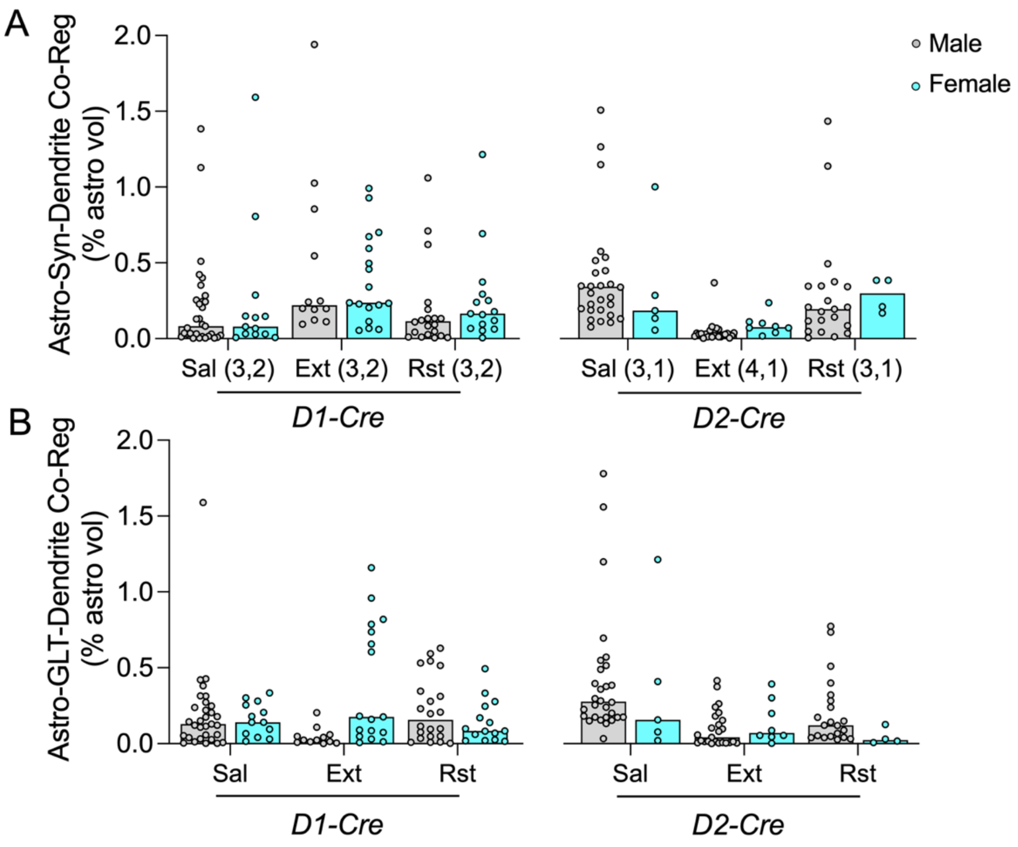


**Fig. S6. Cell-type specific dynamics in astrocyte-synapse and GLT-dendrite association are not sex-dependent.** (**A**) No sex differences were detected in D1- or D2-synaptic co-registration by NAcore astroglia after operant training with heroin (2-way ANOVA $F_{1,196}$=0.7529 p=0.3866). (**B**) Dendritic association of GLT-1 did not differ by sex (2-way ANOVA $F_{1,196}$=0.05632 p=0.8127). Sal, yoked saline; Ext, extinguished; Rst, 15-min cued reinstatement. Animal N shown below bars in (**A**) as (male, female).

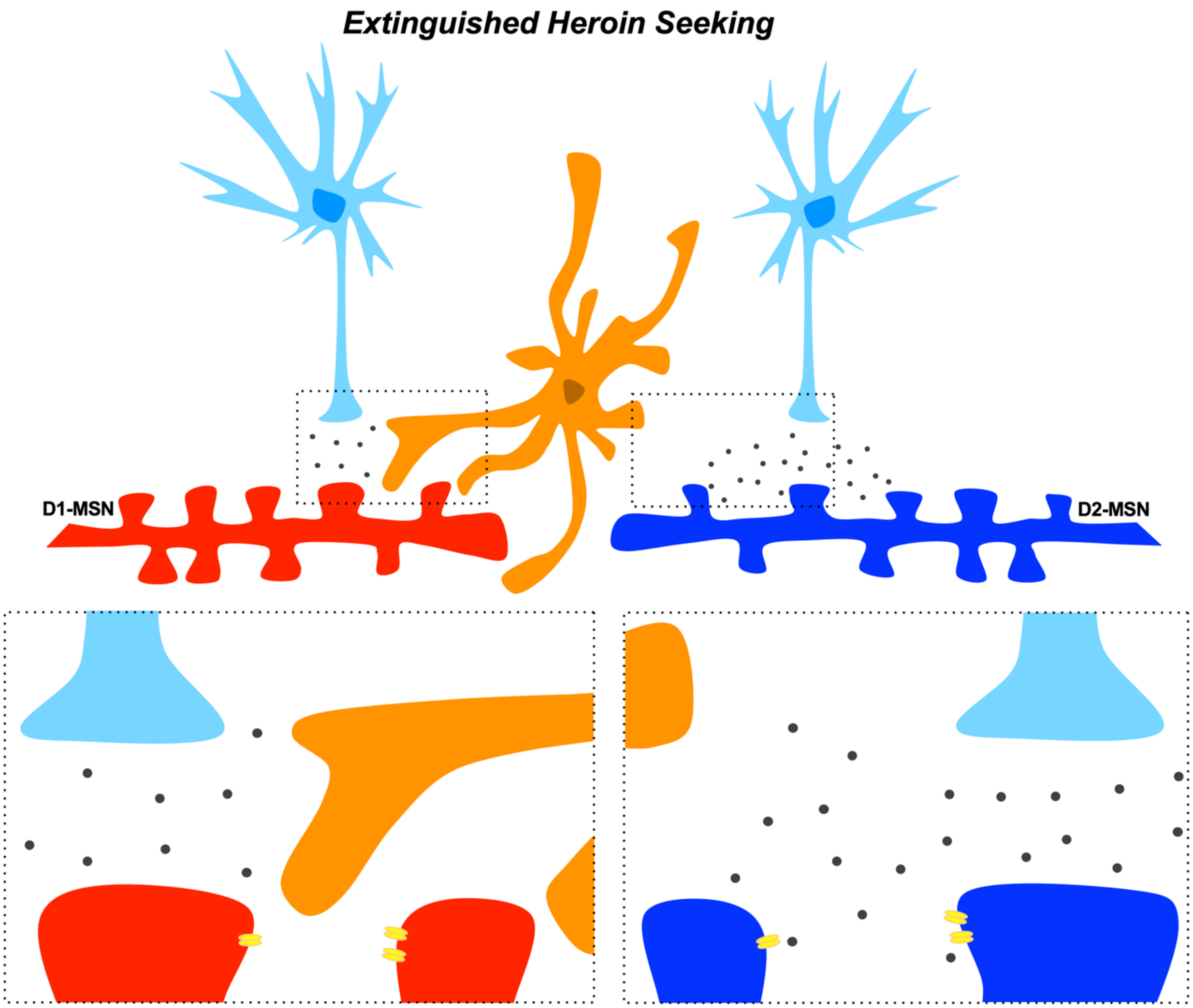


**Fig. S7. Synapse-selective proximity of astrocytes regulates seeking.** Extinguished heroin seeking is characterized by astrocytes exhibiting a high degree of association with synapses from D1-MSNs (left), but retraction from synapses from D2-MSNs (right). High synaptic co-registration by astroglial processes low in GLT-1 (orange) is predicted to induce autoinhibition through spatial buffering of glutamate toward presynaptic inhibitory autoreceptors (*36*). The high co-registration of astroglia with D1-MSN synapses during extinction training is also predicted to shield post-synaptic NR2B receptors (yellow) that produce postsynaptic potentiation when stimulated by glutamate (*33*). These two functions would suppress D1-MSN potentiation during extinction training. Instead, retraction from D2-MSNs after extinction of heroin seeking engages post-synaptic potentiation of D2-MSNs through stimulation of postsynaptic NR2B (*33*), synaptic recruitment (*41*), and loss of autoinhibitory mechanisms at terminals synapsing onto D2-MSNs (*36*). These hypotheses are supported by data showing potentiation of D2-MSNs, but not D2-MSNs during extinction training (*26, 46*). Reversal of this pattern during cued reinstatement would permit potentiation of D1-MSNs and block signaling through D2-MSNs, permitting seeking behavior (*10*).

| D1-Cre | | | | | | D2-Cre | | | | | |
|---|---|---|---|---|---|---|---|---|---|---|---|
| *Saline* | | *Extinguished* | | *Reinstated* | | *Saline* | | *Extinguished* | | *Reinstated* | |
| Astro-Syn-Dendrite Co-Reg (% vol) | Astro-GLT-Dendrite Co-Reg (% vol) | Astro-Syn-Dendrite Co-Reg (% vol) | Astro-GLT-Dendrite Co-Reg (% vol) | Astro-Syn-Dendrite Co-Reg (% vol) | Astro-GLT-Dendrite Co-Reg (% vol) | Astro-Syn-Dendrite Co-Reg (% vol) | Astro-GLT-Dendrite Co-Reg (% vol) | Astro-Syn-Dendrite Co-Reg (% vol) | Astro-GLT-Dendrite Co-Reg (% vol) | Astro-Syn-Dendrite Co-Reg (% vol) | Astro-GLT-Dendrite Co-Reg (% vol) |
| 0.0300 | 0.0076 | 0.7001 | 0.7365 | 0.1932 | 0.2788 | 0.1307 | 0.2724 | 0.0287 | 0.3753 | 0.1782 | 0.1425 |
| 0.2456 | 0.1780 | 0.0605 | 0.1774 | 0.1327 | 0.5443 | 0.1984 | 0.3842 | 0.0155 | 0.1248 | 0.1163 | 0.7739 |
| 0.5098 | 0.2482 | 0.3413 | 1.1587 | 0.1322 | 0.0502 | 0.1127 | 0.1744 | 0.0455 | 0.2622 | 0.3429 | 0.3991 |
| 0.0305 | 0.0043 | 0.6727 | 0.7868 | 0.0913 | 0.2309 | 0.3452 | 0.4887 | 0.3680 | 0.0975 | 0.3757 | 0.1197 |
| 0.2540 | 0.2729 | 0.2288 | 0.6566 | 0.1210 | 0.0947 | 0.2270 | 0.1807 | 0.0200 | 0.1809 | 0.2075 | 0.3220 |
| 0.0708 | 0.0527 | 0.2227 | 0.9594 | 0.1350 | 0.5158 | 0.2557 | 0.2551 | 0.0304 | 0.2420 | 1.4346 | 0.7354 |
| 0.0123 | 0.0277 | 0.0679 | 0.8191 | 0.6213 | 0.3455 | 0.3475 | 0.2966 | 0.0368 | 0.0078 | 0.0115 | 0.5105 |
| 0.0155 | 0.0398 | 0.5954 | 0.6045 | 0.7099 | 0.6298 | 0.1069 | 0.0336 | 0.0127 | 0.1029 | 0.0477 | 0.0883 |
| 0.0144 | 0.0020 | 1.0266 | 0.1080 | 0.0482 | 0.3122 | 0.3589 | 0.1326 | 0.0151 | 0.0145 | 0.3453 | 0.2403 |
| 0.2441 | 0.1499 | 0.1108 | 0.0421 | 0.0080 | 0.0475 | 0.0768 | 0.1742 | 0.0294 | 0.2039 | 1.1386 | 0.1740 |
| 0.2063 | 0.1212 | 1.9399 | 0.0129 | 2.4966 | 0.5931 | 0.1889 | 0.2818 | 0.0124 | 0.4163 | 0.4940 | 0.0402 |
| 0.0212 | 0.0581 | 0.1929 | 0.2035 | 0.0837 | 0.1153 | 1.5075 | 2.9899 | 0.0388 | 0.1537 | 0.0353 | 0.0314 |
| 0.1315 | 0.1291 | 0.2483 | 0.0004 | 0.0262 | 0.0093 | 5.6845 | 1.5592 | 0.0611 | 0.0035 | 0.3478 | 0.0415 |
| 0.0793 | 0.0665 | 0.1987 | 0.0241 | 0.0108 | 0.0082 | 2.0164 | 0.2007 | 0.0687 | 0.0054 | 0.1945 | 0.2818 |
| 0.1483 | 0.1803 | 0.1978 | 0.0131 | 0.0059 | 0.0021 | 0.3009 | 0.1548 | 0.0110 | 0.0584 | 0.0424 | 0.1492 |
| 0.0067 | 0.0151 | 0.2338 | 0.1819 | 0.0194 | 0.0102 | 1.1487 | 0.3968 | 0.0391 | 0.0448 | 0.0847 | 0.0475 |
| 0.0293 | 0.0416 | 0.1124 | 0.1596 | 0.0432 | 0.0984 | 0.4489 | 0.5708 | 0.0705 | 0.0551 | 0.2491 | 0.0681 |
| 0.1392 | 0.1404 | 0.9283 | 0.0909 | 0.0108 | 0.0757 | 0.3398 | 0.1734 | 0.1116 | 0.3930 | 0.2068 | 0.0527 |
| 0.0670 | 0.0305 | 0.0538 | 0.0847 | 0.1098 | 0.1977 | 0.5152 | 1.1983 | 0.0172 | 0.0019 | 0.1428 | 0.0393 |
| 0.4016 | 0.2204 | 0.2374 | 0.0477 | 3.9462 | 0.2067 | 0.2718 | 0.3590 | 0.0710 | 0.1988 | 0.1213 | 0.1213 |
| 0.2838 | 0.4279 | 0.9920 | 0.0307 | 0.2386 | 0.0353 | 0.2250 | 0.1535 | 0.1009 | 0.0544 | 0.0052 | 0.0276 |
| 1.1285 | 0.3126 | 0.4975 | 0.0118 | 1.0601 | 0.5309 | 1.2654 | 0.5146 | 0.2368 | 0.0877 | 0.3853 | 0.0165 |
| 0.3527 | 0.2036 | 0.2042 | 0.0075 | 0.0956 | 0.1388 | 2.2984 | 1.7794 | 0.0798 | 0.3014 | 0.3851 | 0.1269 |
| 0.0352 | 0.1150 | 0.4574 | 0.0740 | 0.3731 | 0.2772 | 0.5759 | 0.1510 | 0.0544 | 0.0044 | 0.1684 | 0.0301 |
| 0.0048 | 0.1748 | 0.5457 | 0.0033 | 0.2509 | 0.2463 | 0.1536 | 0.5491 | 0.0236 | 0.0132 | 0.2111 | 0.0072 |
| 0.0941 | 0.3145 | 0.2411 | 0.0600 | 0.2393 | 0.0616 | 0.1875 | 0.2406 | 0.0797 | 0.0416 | | |
| 0.2877 | 0.2814 | 0.1222 | 0.0089 | 0.1581 | 0.0183 | 0.3488 | 0.2252 | 0.0117 | 0.0136 | | |
| 0.0284 | 0.2051 | 0.8550 | 0.0477 | 0.0052 | 0.0247 | 0.4335 | 0.3767 | 0.0328 | 0.0070 | | |
| 1.5922 | 0.3352 | 0.0949 | 0.0229 | 0.6927 | 0.4945 | 0.5357 | 0.6957 | 0.0184 | 0.0157 | | |
| 0.8063 | 0.2548 | | | 0.1775 | 0.1694 | 0.1224 | 0.1730 | 0.0357 | 0.0801 | | |
| 0.1473 | 0.3016 | | | 1.2145 | 0.3333 | 0.2860 | 1.2131 | 0.0293 | 0.0202 | | |
| 0.0076 | 0.0951 | | | 0.0673 | 0.0841 | 0.1845 | 0.4097 | 0.0047 | 0.0005 | | |
| 0.0311 | 0.1405 | | | 0.1652 | 0.0209 | 0.0565 | 0.0780 | 0.0023 | 0.0039 | | |
| 0.4217 | 1.5885 | | | 0.0536 | 0.0818 | 0.1340 | 0.0199 | | | | |
| 1.3833 | 0.3813 | | | 0.1204 | 0.0696 | 1.0013 | 0.1578 | | | | |
| 0.2289 | 0.4215 | | | 0.0628 | 0.0153 | | | | | | |
| 0.1299 | 0.1208 | | | 0.2941 | 0.0977 | | | | | | |
| 0.0039 | 0.1085 | | | | | | | | | | |
| 0.0373 | 0.1434 | | | | | | | | | | |
| 0.0312 | 0.0317 | | | | | | | | | | |
| 0.0822 | 0.2388 | | | | | | | | | | |
| 0.0097 | 0.0025 | | | | | | | | | | |
| 0.0040 | 0.0594 | | | | | | | | | | |
| 0.0021 | 0.0123 | | | | | | | | | | |

**Table S1. Dendrite-specific co-registration values.** Triple co-registration of astroglia with Synapsin I and labeled dendrites, as well as triple co-registration of astroglia with GLT-1 and labeled dendrites from D1- and D2-Cre rats are presented as % astrocyte volume.
